# Deep metagenomics examines the oral microbiome during dental caries, revealing novel taxa and co-occurrences with host molecules

**DOI:** 10.1101/804443

**Authors:** J.L. Baker, J.T. Morton, M. Dinis, R. Alverez, N.C. Tran, R. Knight, A. Edlund

## Abstract

Dental caries is the most common chronic infectious disease globally. The microbial communities associated with caries have mainly been examined using relatively low-resolution 16S rRNA gene amplicon sequencing and/or using downstream analyses that are unsound for the compositional nature of the data provided by sequencing. Additionally, the relationship between caries, oral microbiome composition, and host immunological markers has not been explored. In this study, the oral microbiome and a panel of 38 host markers was analyzed across the saliva from 23 children with dentin caries and 24 children with healthy dentition. Metagenomic sequencing, followed by investigation using tools designed to be robust for compositional data, illustrated that several *Prevotella* spp. were prevalent in caries, while *Rothia* spp. were associated with the health. The contributional diversity (extent to which multiple taxa contribute to each pathway) of functional pathways present in the oral microbiome was decreased in the caries group. This decrease was especially noticeable in several pathways known to impede caries pathogenesis, including arginine and branched-chain amino acid biosynthesis. 10 host immunological markers were found to be significantly elevated in the saliva of the caries group, and microbe-metabolite co-occurrence analysis provided an atlas of relationships contributing to the bi-directional influence between the oral microbiome and the host immune system. Finally, 527 metagenome-assembled genomes were obtained from the metagenomics data, representing 151 species. 23 taxa were novel genera/species and a further 20 taxa were novel species. This study thus serves as a model analysis pipeline that will tremendously expand our knowledge of the oral microbiome and its relationship to dental caries once applied to large populations.

## Introduction

Dental caries is the most common chronic infectious disease and will afflict well over half of the global human population at some point in their lives. Dental caries is particularly problematic in children, where it is five times more common than asthma, the second most common chronic disease. This extreme prevalence translates to an extraordinary economic burden. Caries disproportionally afflicts vulnerable populations least able to access and afford proper treatment (1). Historically, members of the acid-producing and acid-tolerant mutans group of streptococci, particularly the paradigm species of the group, *Streptococcus mutans*, were considered the etiologic agents of the disease (2). While caries is certainly an infectious and transmissible disease caused by the oral microbiota, it is now understood to be multifactorial and ecology-based, because although mutans streptococci are commonly associated with caries, they are neither necessary nor sufficient to cause disease (3). Interplay between host genetic and immunological factors, diet and hygiene habits, and the oral microbiota affect clinical development of the disease (1, 4).

Second-generation sequencing techniques enabled many studies characterizing the oral microbiome to distinguish healthy individuals from those with dental caries. The highlights and challenges of this progress have been the subject of several excellent recent reviews (5–7). These studies of the oral microbiome in the context of dental caries have yielded varied results, particularly vis-à-vis the association (and therefore inferred importance) of *S. mutans* and other taxa with the disease. This has rightfully led to some debate regarding the long-standing dogma that *S. mutans* is a keystone species in caries pathogenesis (3, 8, 9). These studies used widely different sampling techniques, library prep methods, and data analysis methods, which contribute substantially to variation among studies. Furthermore, ethnicity, immunology, diet, hygiene and other factors also likely contribute variability and are difficult to control. Finally, because microbiome sequencing provides data in the form of relative abundances, inferring absolute fold-changes or correlations is inherently problematic (10–13). Numerous microbiome studies have drawn biological conclusions based on the application of conventional statistical tools to compositional data, which has been shown to have unacceptably high false discovery rates and lead to spurious hypotheses (11).

The overwhelming majority of studies examining the microbiome associated with caries have utilized 16S rRNA gene amplicon sequencing (“16S sequencing”) (9). While 16S sequencing is widely used, relatively inexpensive, and has significantly advanced the field of microbiology, there are a number of disadvantages to using this technique. These include the biases introduced during the PCR amplification step (i.e. different 16S amplicons amplify at different efficiencies for various species and genera), biases due to the fact that many taxa encode differing copy numbers of the 16S gene, and the inability to distinguish organisms at the strain, species, or, occasionally, even higher taxonomic level (14–17). For example, *S. mutans,* an overtly cariogenic species, and *Streptococcus gordonii,* largely associated with good dental health, are both simply identified as ‘*Streptococcus’* in many 16S-based studies. In addition, studies have suggested that compositional differences at the species level can, at times, be less reflective of the health of a microbial community than differences in the metabolic functions of the community (i.e. interpersonal microbial taxonomic profiles may be significantly different, but communities remain more or less functionally equivalent) (18). Although bioinformatics tools, such as PICRUSt (19), predict community functions based on 16S amplicon sequencing, evidence continues to mount that the pan-genomic (i.e. intra-species) diversity of many individual taxa is massive, limiting the utility of such predictions when the key genes involved in a biological process are not conserved phylogenetically or among strains of a species. In the case of dental caries, this strain-to-strain variation is well-documented to affect both virulence of pathogens (e.g. *S. mutans*) and protective effects of commensals (e.g. *S. gordonii*), further illustrating a need for studies utilizing more in-depth sequencing and analysis methods (5). Although several shotgun metagenomic surveys of the oral microbiota have been performed (20–23), they have not employed differential ranking techniques with consistent reference frames, as detailed in Morton et al. 2019 (13).

In this study, shotgun metagenomic sequencing was used to examine the oral microbiome of 23 children with severe dentin caries, compared to 24 children with good dental health. The study groups were not equal numbers due to difficulties in recruitment. This shotgun sequencing, followed by investigation using state-of-the-art tools for analysis of compositional data and generation of metagenome-assembled genomes (MAGs), allowed identification and relative quantification of species and strain-level taxa, analysis of the functional pathways present, and reconstruction of high-quality species-level genomes, including those of nearly 50 novel taxa.

The oral samples utilized for metagenomic sequencing were also tested for the presence of 38 salivary immunological markers, 10 of which were significantly elevated in the caries group. Although dental caries pathogenesis originates on the non-shedding, hard surface of the tooth, the immunology of the human host nevertheless plays a critical role in disease prevention or progression. Both the innate and adaptive arms of the immune system influence caries, and past proof-of-principle studies explored the possibility of several active and passive vaccine strategies to prevent caries (reviewed in (24, 25)). In recent years, however, research regarding the immunological component of dental caries has lagged well behind study of the microbiological component. This is due at least in part to the prevailing perception of dental caries as an ‘over-the-counter’ disease (26). The oral microbiota clearly influences the host immune system and vice-versa, and it is likely that in cases of good oral health, the immune system has evolved to tolerate and facilitate maintenance of a commensal, yet territorial oral microbiota that prevents the establishment of foreign pathogens (27). The co-occurrence analysis performed in this study between microbial features and salivary immunological markers provides the first detailed look at potential cross-talk between the oral microbiota and the immune system of the human host during advanced dental caries compared to health.

## RESULTS

### Study design (Figure 1A)

Details of the clinical sampling, as well as inclusion and exclusion criteria are provided in the MATERIALS AND METHODS section. In brief, 47 participants aged 4-11 received a comprehensive oral examination, and their dental caries status was recorded using decayed (d), missing due to decay (m), or filled (f) teeth in primary and permanent dentitions (dmft/DMFT) by a clinician (Fig 1A). A summary of the collected subject metadata is provided in Table S1. Subjects were dichotomized into two groups: healthy (0 decayed, missing, or filled teeth [DMFT]), or caries (*≥* 2 active caries lesions with penetration through the enamel into the underlying dentin, only lesions at least 2 mm in depth were considered). All subjects provided 2 ml of unstimulated saliva and 2ml of stimulated saliva, which was clarified by centrifugation. DNA was extracted from the stimulated saliva samples and subjected to Illumina sequencing and metagenomics analysis as described in MATERIALS AND METHODS. The concentration of 38 immunological markers in the unstimulated saliva samples were determined using a multiplex Luminex bead immunoassay performed by Westcoast Biosciences. Dataset S1 contains the comprehensive output of the Luminex assay. A detailed summary of the bioinformatics tools and pipelines used in this study are provided in Figure 1B.

**Fig 1.**
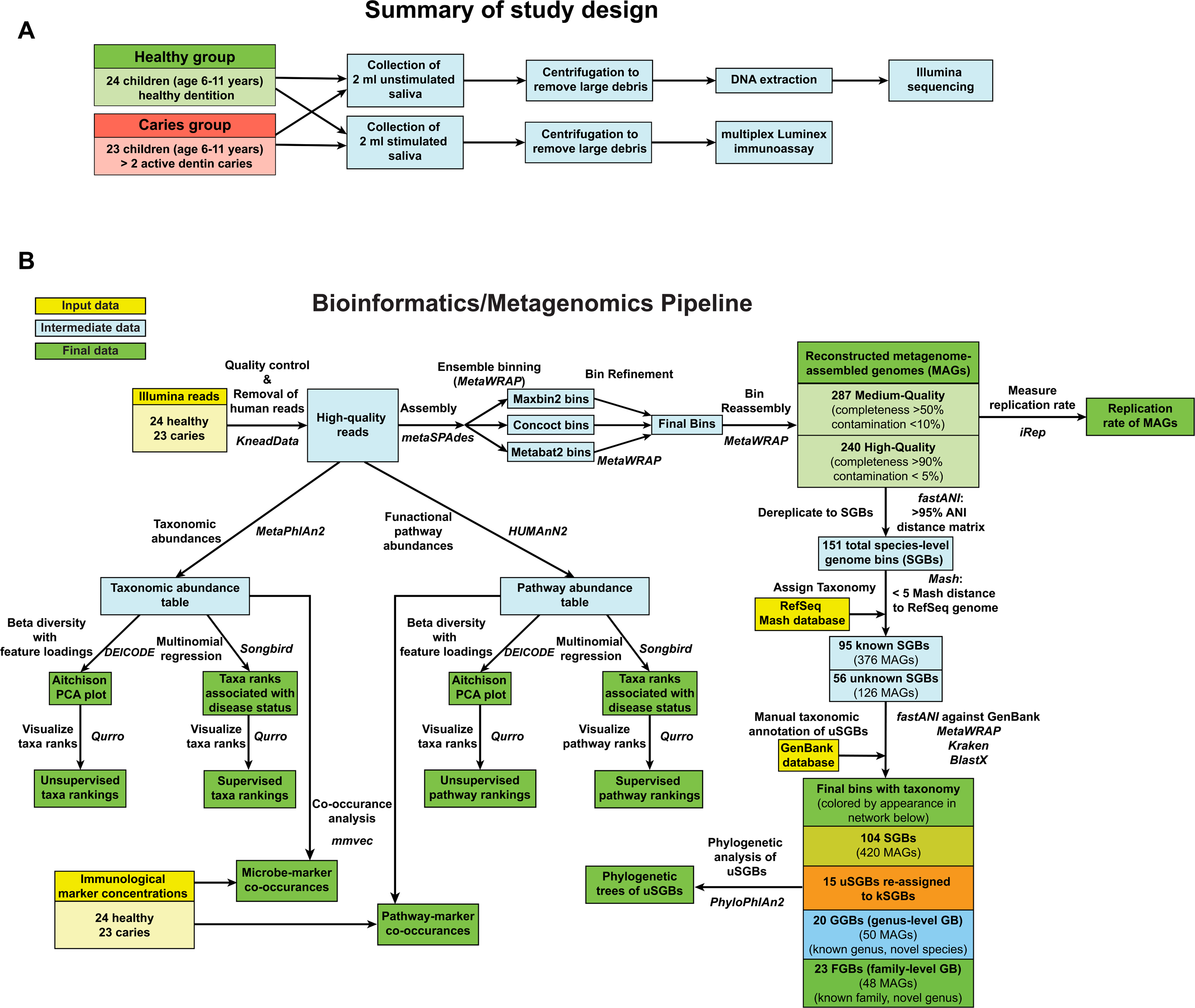
Overview of study design and bioinformatics methods. **(A)** Flow chart illustrating the steps taken to get from clinical specimen to bioinformatics data. **(B)** Flow chart illustrating the computational methodology utilized in this study. Input data is in yellow boxes, intermediate data is in blue boxes, and final data is in green boxes. For each step, the tool(s) or package(s) used are provided in italics. The ‘Final bins with taxonomy’ box is color-coded to match the metagenome-assembled genome (MAG) average nucleotide (ANI) network in Figure 5.

### In this study group, *Prevotella* spp. were associated with disease, while *Rothia* spp. were associated with good dental health

Following quality control performed by KneadData (available at https://bitbucket.org/biobakery/kneaddata), MetaPhlAn2 (28) was utilized to determine the relative abundance of microbial taxa within each sample. The most abundant taxa across all samples belonged to the taxonomic groups *Prevotella, Veillonella, Porphyromonas, Rothia, Haemophilus, Streptococcus, and Saccharibacteria* (Figure 2A and S1). Notable trends included a higher relative abundance of *Rothia* spp*., Porphyromonas* sp*. oral taxon 270, Haemophilus parainfluenzae* and *Streptococcus sanguinis* in the saliva from healthy children (Figure 2A). Meanwhile, although *S. mutans* was detected at higher relative abundances in the saliva derived from the children with caries, *S. mutans* and the other canonical caries pathogen, *Streptococcus sobrinus,* were observed in comparably low relative abundances overall, and only within 11 and 3 samples, respectively (*S. mutans*: 7 caries and 4 healthy; *S. sobrinus:* 2 caries, 1 healthy). The complete taxonomic table generated by MetaPhlAn2 is provided in Table S2 and a heatmap is provided in Figure S1.

**Fig 2.**
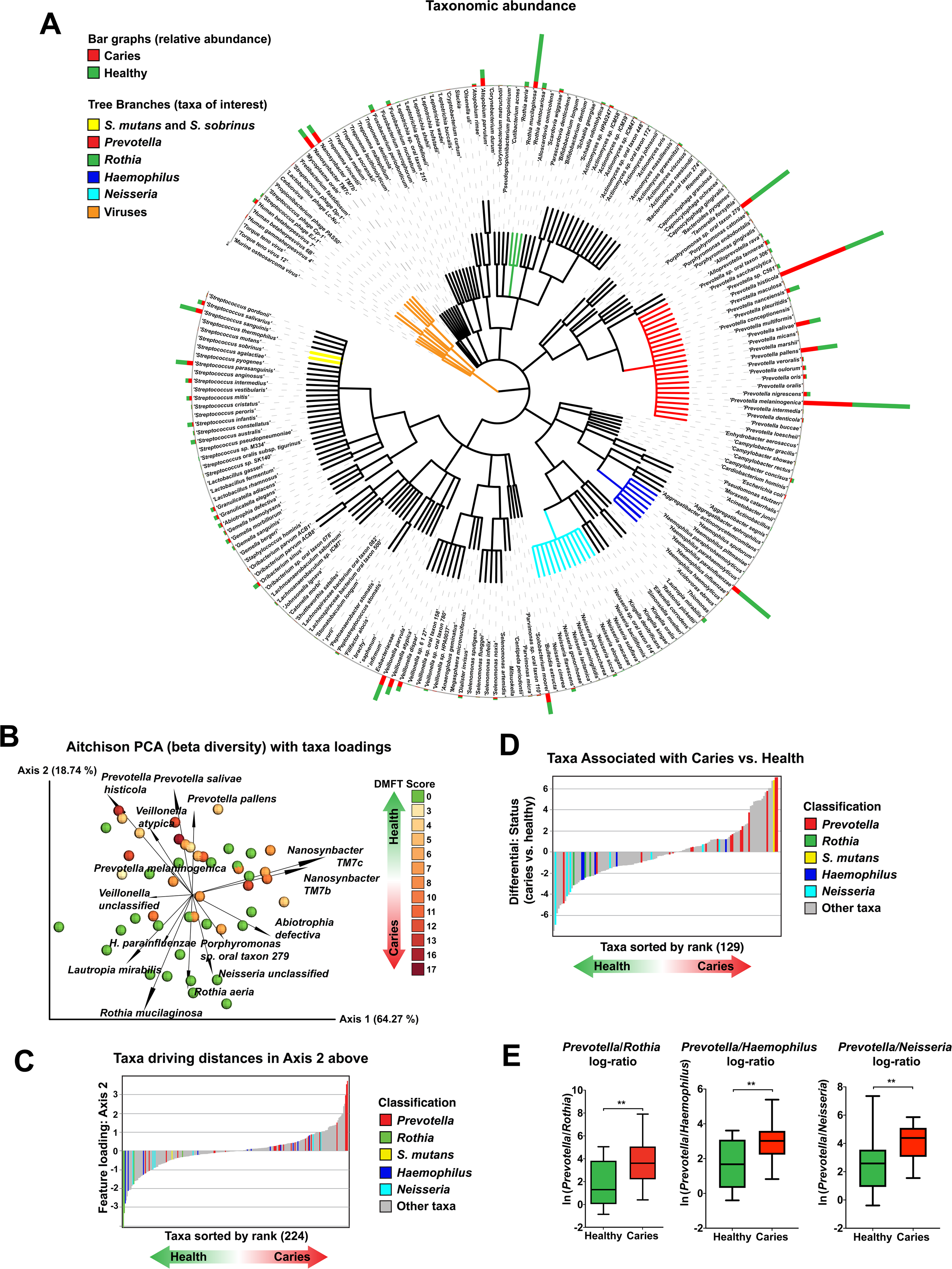
Significant taxonomic differences in the oral metagenome between healthy children and children with caries. **(A) Species abundance.** Phylogenetic tree illustrating the species present across the saliva metagenomes. The relative abundance of each taxa is represented by the bar graph at the end of each leaf, with the relative abundance in the healthy group in blue and the caries group in red. Taxa of interest are highlighted with colored leaves on the tree: *Streptococcus mutans* and *Streptococcus sobrinus* = yellow; *Prevotella* spp.= red; and *Rothia* spp. = green. **(B) Beta Diversity.** Compositional biplot visualized in Emperor (39) and generated using DEICODE (robust Aitchison PCA)(30). Data points represent individual subjects and are colored with a gradient to visualize DMFT score, indicating severity of dental caries. Feature loadings (i.e. taxa driving differences in ordination space) are illustrated by the vectors, which are labeled with the cognate species name. **(C) Ranking of RPCA Axis 2 feature loadings.** Qurro-produced bar chart illustrating the sorted ranks of the feature loadings of Axis 2 from Figure 1B, corresponding to the main RPCA space separation between the healthy and caries groups. *Prevotella* spp. are highlighted in red, while *Rothia* spp. are highlighted in green, *S. mutans* is highlighted in yellow, *Haemophilus* spp. are highlighted in dark blue, and *Neisseria* spp. are highlighted in light blue. **(D) Differential rankings of taxa associated with disease status.** Qurro-produced bar chart illustrating the sorted differential rankings of taxa associated with disease status determined by Songbird (13). *Prevotella* spp. are highlighted in red, *Rothia* spp. are highlighted in green, and *S. mutans* is highlighted in yellow, *Haemophilus* spp. are highlighted in dark blue, and *Neisseria* spp. are highlighted in light blue. (E) The log-ratio of *Prevotella* spp*./Rothia* spp. is significantly increased in caries. Bar chart illustrating the log2 ratio of *Prevotella* spp*./Rothia* spp. across the healthy and caries sample groups. **, denotes statistical significance, based on a Welch’s *t*-test (p = 0.001). No samples were dropped as all samples contained these 4 taxa.

Alpha diversity (within-sample diversity), was calculated using QIIME2 (29), was not significantly different between the healthy and caries groups, and was not correlated to DMFT scores, Age, Lesion Depth, or the Number of Lesions, according to the Shannon and Simpson metrics (data not shown). Beta diversity (between sample diversity) was determined using DEICODE (30), which utilizes matrix completion and robust Aitchison principal component analysis (RPCA), providing several advantages over other tools, including the ability to accurately handle sparse datasets (e.g. in most microbial communities, most taxa are not present in a majority of samples), scale invariance (negating the need for rarefaction) and preservation of feature loadings (facilitating the analysis of which taxa are driving the differences in the ordination space)(30). DEICODE illustrated a clear difference in beta diversity between the healthy and caries subject groups (Figure 2B), which was statistically significant based upon a PERMANOVA (p = 0.003), and occurred mainly along Axis 2 (the vertical axis). The 15 species that were the most significant drivers of distance in ordination space are illustrated by the vectors in Figure 2B. Qurro (doi:10.5281/zenodo.3369454) was used to visualize and identify taxa driving the differences along Axis 2, which seemed to correspond to separation between the healthy and caries samples in the ordination space (Figure 2C). *Prevotella histicola, Prevotella salivae,* and *Prevotella pallens* were the top 3 drivers in the positive direction along Axis 2 (corresponding to the caries samples), while *Rothia mucilaginosa* and *Rothia aeria* were the top 2 drivers in the negative direction along Axis 2 (corresponding to the healthy samples) (Figure 2C). Certain *Neisseria* and *Haemophilus* spp. also seemed to be generally associated with the healthy samples (Figure 2B and C). *S. mutans,* the classic caries pathogen, did not appear to be a major driver of beta diversity, according to DEICODE (Figure 2C).

### Differential ranking reveals *S. mutans* is significantly associated with caries

As the patterns observed above used taxa ranked in an unsupervised manner, it was important to determine which taxa were directly associated with disease status (and not simply Axis 2 of the DEICODE biplot). Songbird (13) and Qurro were used to respectively calculate and display the differential ranks of taxa directly associated with health versus disease (Figure 2D). Because this method is sensitive to sparsity, only species observed in at least 10 samples and having over 10,000 total predicted counts were analyzed here. A similar pattern was observed using this approach, with several *Prevotella* species the most correlated taxa the caries samples and all 3 observed *Rothia* species correlated with the health samples, as were most *Neisseria* and *Haemophilus* spp. (Figure 2D). Differential ranks are listed in Table S3. Interestingly, *S. mutans,* the classic caries pathogen, was the third-highest-ranked taxon in association with the caries samples (Figure 2D). Log ratios are a preferable way to examine differences within compositional datasets (13), and as *Prevotella* spp. were significantly associated with the caries samples and *Rothia, Haemophilus* and *Neisseria* spp. were strongly associated with the healthy samples, according to both unsupervised and supervised methods, the log ratios of *Prevotella* to *Rothia, Haemophilus* and *Neisseria* were examined. In all three cases, the log-ratio with *Prevotella* as the numerator was significantly higher in the caries group compared to the healthy group, indicating that the ratio of these taxa may have clinical significance and be a useful marker of disease (Figure 2E). Although the ranking of *Prevotella, Rothia, Haemophilus,* and *Neisseria* in regards to disease status is generally concordant between DEICODE and Songbird, there is some discrepancy in the ranks of other taxa, mainly low-abundance. This is likely due the nature of multinomial regression (employed by Songbird), in which features with low counts can have a larger fold change than features with high counts. *Rothia* is a genus that has received little attention and has been previously associated with either caries or good dental health, depending on the study (possibly due to use of non-compositional data analyses), indicating that this taxon demands further examination (13, 31, 32).

### Fungi and viruses are present in low numbers in the oral microbiomes examined in this study

Unlike 16S sequencing, metagenomic sequencing detects viruses and eukaryotes in addition to bacteria. 12 viruses were detected in this study, including several human herpesviruses and several bacteriophage (Figure 2A). The viruses were detected at relatively low frequency and did not appear to be significant drivers of beta diversity in this study group (Figure 2). The fungal pathogen, *Candida albicans* is known to be involved in pathogenesis in many cases of dental caries (reviewed in (33)), therefore it was surprising that it was not detected by MetaPhlAn2 in this study. Mapping Illumina reads directly to the *C. albicans* genome indicated the presence of *C. albicans* in the samples, but the number of reads was small, and thus any fungal pathogens present in the study group were likely to be below the threshold of detection employed in taxonomic quantification by MetaPhlAn2 (data not shown). It is possible that the extraction methods used did not efficiently lyse fungal cells, leading to an observed underrepresentation.

### There is a decrease in the diversity of functional pathways in the oral microbiome of children with caries

To examine the differential representation of particular microbial metabolic and biosynthetic pathways in caries and health, HUMAnN2 (34) analysis was performed on the quality-controlled Illumina reads. The resulting pathway abundance table (Table S4, Figure S2) was analyzed using QIIME2, DEICODE, and Songbird. The functional pathways of the oral microbiome from the caries group had a lower alpha diversity, as measured using the Shannon Index (Figure 3A), and this metric inversely correlated with the number of lesions (Figure S3). 99.97% of the functional pathway beta diversity was explained by one PCA axis in the DEICODE biplot (Figure 3B), and disease status was less associated with functional than taxonomic pathway beta diversity (PERMANOVA, P = 0.016 vs 0.003). This reduction in variance made it difficult to interpret whether any pathways were correlated to disease status. Rare, low abundance features were most strongly associated with the caries group, while several pathways, including anaerobic and aerobic energy metabolism, were associated with health as shown in results from DEICODE (Figure 3B) and Songbird (Table S5). One of the advantages of HUMAnN2 is the ability to stratify pathways by taxa and examine contributional diversity, the extent to which multiple taxa contribute to particular functional pathway (34). Across the 69 core functional pathways which were present in all 47 samples and had more than 3 contributing taxa, contributional diversity was decreased in the caries group compared to the healthy group (Figure 3C). This was more noticeable along the x-axis, corresponding to alpha diversity, and congruent with the data provided in Figures 3A and B, which was agnostic to contributing taxa. Contributional diversity was examined for several specific pathways identified as health-associated in Songbird, and several pathways that have a well-established connection to the prevention of caries pathogenesis, including arginine biosynthesis (35), unsaturated fatty acid biosynthesis (36), branched-chain amino acid (BCAA) biosynthesis (37), and urea metabolism (38). The differences seen in several of these pathways were particularly striking (Figures 3D-H). Overall, the data from the functional analyses of the oral microbiome indicates that there is less variation between caries and health in terms of functional pathways compared to taxa. However, several pathways were clearly associated with the healthy samples, including several where the physiological relationship to caries pathogenesis is understood.

**Fig 3.**
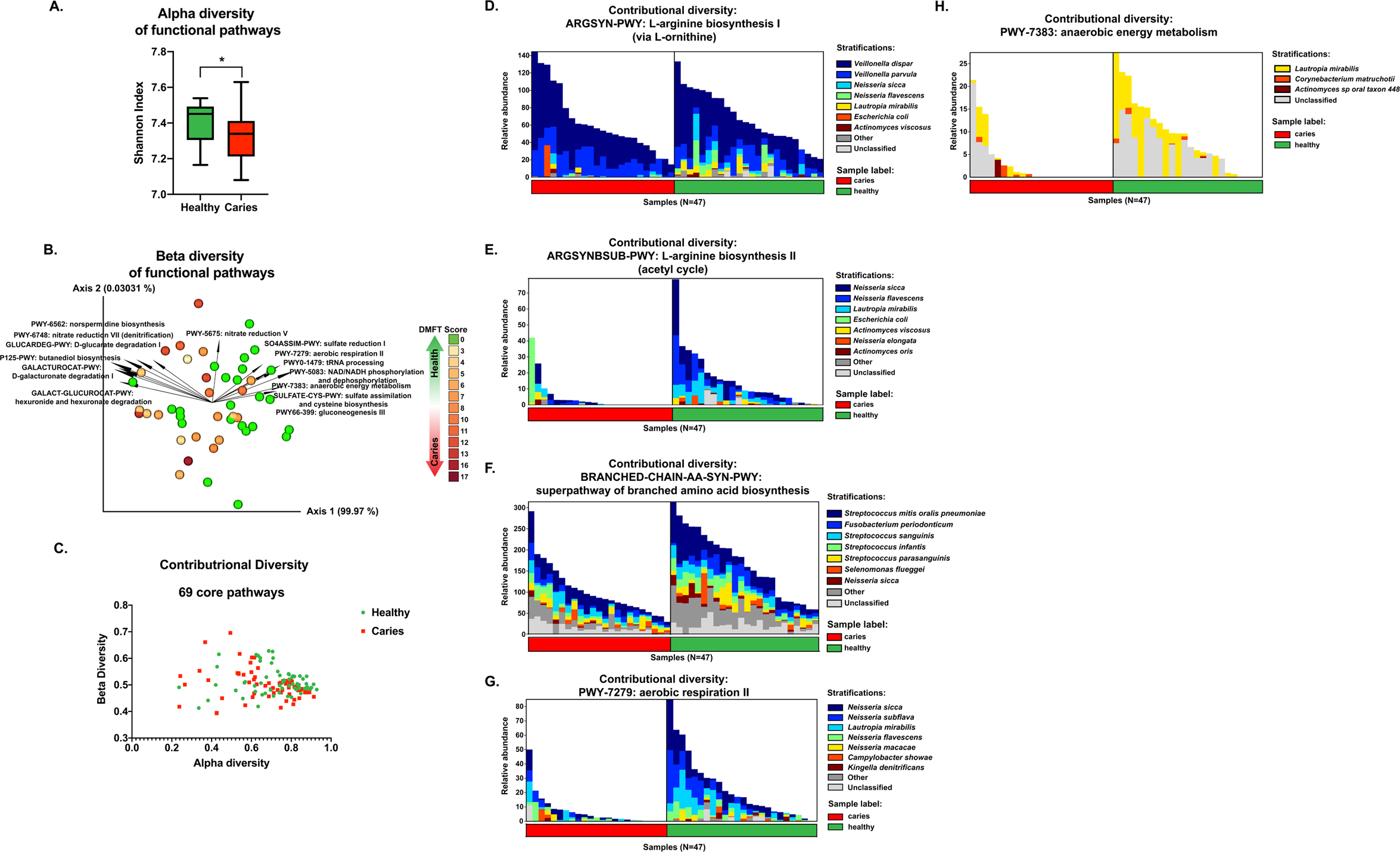
Profiling of functional pathways illustrates differences between health- and caries-associated oral microbiota. **(A) A greater diversity of functional pathways are present in the healthy group.** Bar chart illustrating the alpha diversity (Shannon Index) of the functional pathways present in the healthy and caries groups, as determined by HUMAnN2 (34) analysis. *, indicates statistical significance, based upon a Kruskal-Wallis test (p = 0.0136). **(B) Beta diversity of functional pathways.** 3D PCA plot generated using DEICODE (robust Aitchison PCA)(30). Data points represent individual subjects and are colored with a gradient to visualize DMFT score, indicating severity of dental caries. Feature loadings (i.e. functional pathways driving differences in ordination space) are illustrated by the vectors, which are labeled with the cognate pathway name. **(C) Contributional diversity of 69 core pathways.** Scatter plot indicating alpha and beta diversities of 69 functional pathways which were found across all samples. **(D-H) Contributional diversity of pathways of interest to caries pathogenesis.** Stacked bar charts illustrating the relative abundance and contributional diversity of the indicated pathways across the samples (D, L-arginine biosynthesis I; E, L-arginine biosynthesis II; F, branched chain amino acid biosynthesis).

### 10host salivary immunological markers are more abundant in the saliva of children with caries than children with good dental health, and co-occur with *Prevotella histicola, Prevotella salivae,* and *Veilonella atypica*

To investigate differences in the oral immunological profile of healthy children compared to children with caries, a Luminex bead assay was used to quantify 38 known immunological markers. Of these 38 molecules, 7 were at undetectable levels in >50% of the samples (the columns on the far right of Table S1, with a grey background), and thus were not analyzed further. Based on a Welch’s t-test, 10 of the remaining salivary immunological markers were found at significantly higher concentrations in the saliva of children with caries. These were: epidermal growth factor (EGF), interleukin-10 (IL-10), granulocyte-colony stimulating factor (G-CSF), interleukin-1 receptor agonist (IL1-RA), granulocyte/macrophage-colony stimulating factor (GM-CSF), macrophage-derived chemokine (MDC), interleukin-13 (IL-13), interleukin-15 (IL-15), and interleukin-6 (IL-6) (Figure 4A-J). None of the immunological markers were significantly correlated with alpha diversity of the taxa or functional pathways within the samples (data not shown). To examine co-occurrences between specific bacterial species and immunological markers, MMvec (https://github.com/biocore/mmvec) was used to create microbe-metabolite vectors, which were visualized using the QIIME2 Emperor plugin (39). Interestingly, there was a noticeable separation of the directionality of several vectors representing taxa associated with caries (e.g. *Prevotella histicola, Prevotella salivae,* and *Veillonella atypica*) (Figure 4K). These vectors indicated co-occurrence with EGF, IL-10, and IL-1RA (Figure 4K). *Rothia mucilaginosa, Haemophilus parainfluenzae, Streptococcus australis,* and unclassified *Neisseria* formed a cluster of health-associated vectors, and displayed co-occurrence with MCP3 and VEGF (Figure 4K). GRO, MIP-1b, IP-10, MIP-1a, and IL_8 did not appear to have a high co-occurrence with any taxa (Figure 4K). Interestingly, although Streptococcal species did not appear to be large drivers of beta diversity (Figure 2B), a number of Streptococcus species did have considerable co-occurrence with host immunological markers (Figure 4K). A similar approach was attempted to examine co-occurrences between functional pathways and the host markers, but MMvec was developed for taxa-metabolite co-occurrences, and the low dimensionality of the functional pathway data made interpreting the results difficult (Figure S4).

**Fig 4.**
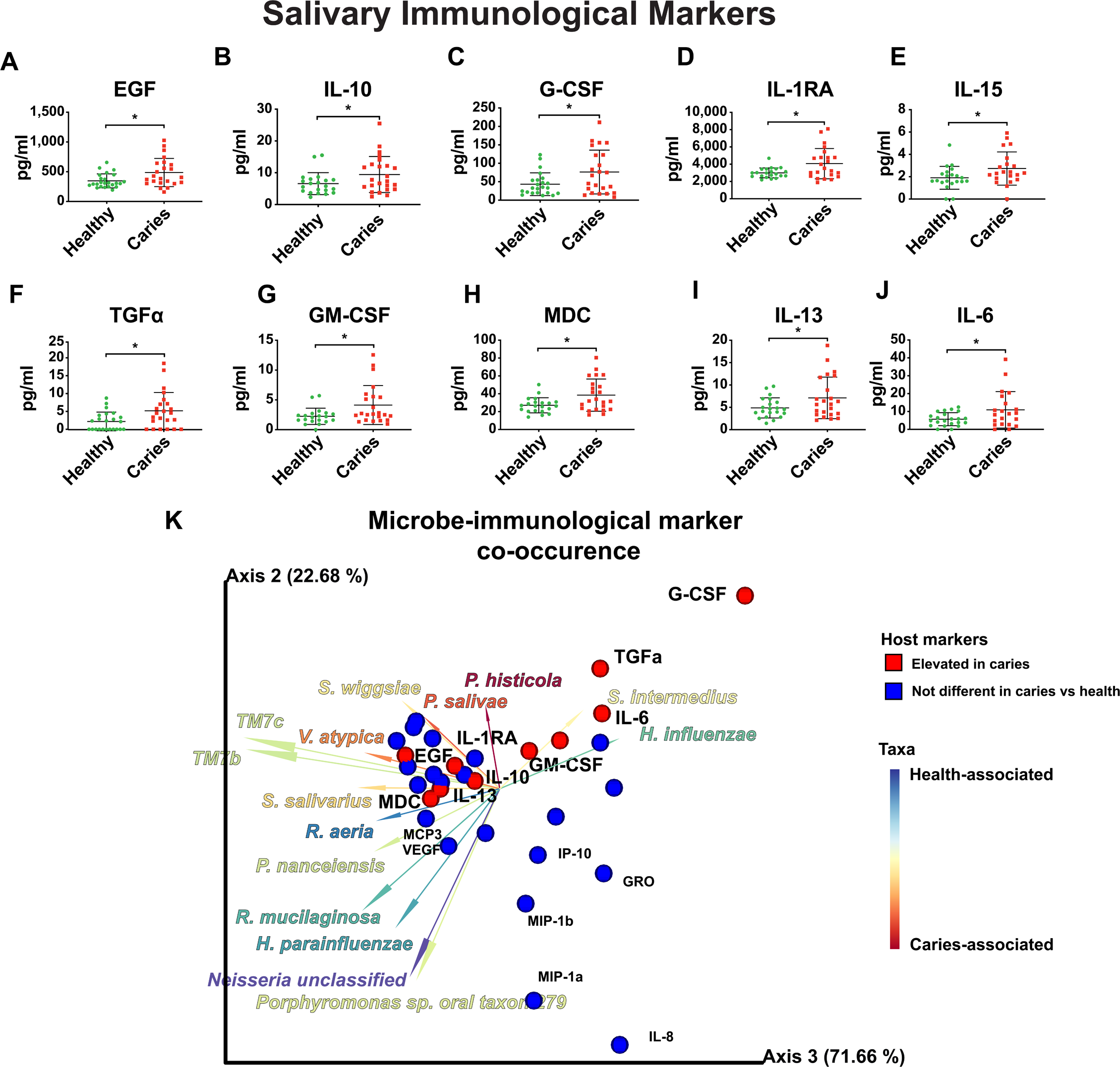
Significant differences in the salivary immunological profile of healthy children and children with caries. **(A-J)** Swarm plots illustrating the 10 immunological markers, **(A)** EGF**, (B)** IL-10, **(C)** G-CSF, and **(D)** IL1-RA, **(E)** IL-15, **(F)** TGF*α*, **(G)** GM-CSF, **(H)** MDC, **(I)** IL-13, and **(J)** IL-6, which were significantly different between healthy and caries subject groups. *, p < 0.05, based on a Welch’s t-test. **(K) Microbe-immune marker co-occurence.** Biplot illustrating the co-occurrence of oral taxa with immune markers. The 31 detected immune markers are represented by spheres, while 15 bacterial taxa with high differential ranks are represented by vectors. Red spheres indicate host markers that were elevated in caries, while blue spheres indicate host markers that were not significantly different between caries and health. Vectors are colored by Songbird ranks (Figure 2D) indicating association with caries versus health. Several immunological markers of interest are labeled.

### Assembly of metagenome assembled genomes (MAGs) recovers 527 medium and high-quality genomes, 20 representing novel species and 23 representing novel genera and species

A more detailed description of the MAG recovery and results is provided in Supplemental Note 1. The metagenomics pipeline illustrated in Figure 1B yielded 527 bins that were of at least Medium Quality according to the guidelines set forth by the Genomic Standards Consortium (GSC) regarding the Minimum Information about a Metagenome-Assembled Genome (MIMAG) (>50% completeness, <10% contamination) (Table S6) (40). A separate assembly was performed for each sample, as opposed to a co-assembly of all samples, an alternative approach used by some studies. The pros and cons of a co-assembly versus individual assemblies have been discussed previously (41). As a result of the individual assemblies, many of the 527 MAGs were likely to represent redundant species across samples. Following dereplication using fastANI (42) and taxonomic assignment using Mash (43), there were 95 known species level genome bins (kSGBs), representing 376 MAGs and 56 unknown SGBs (uSGBs), representing 126 MAGs (Figure 5A). Further examination of the uSGBs reassigned 15 uSGBs, representing 30 MAGs, to kSGBs, as they had >95% ANI match in GenBank (Table S7). 20 uSGBs, representing 50 MAGs, that had 85%-95% ANI match to a GenBank genome were termed genus-level genome bins (GGBs), as the genus can be assigned with a fair amount of confidence, while the species appears to be not previously described. 23 bins, representing 48 MAGs had no match reference in GenBank with an ANI >85%. These were termed family-level genome bins (FGBs), as the family or higher-level taxa can be inferred, but the MAGs likely represent novel genera. Although the GGBs and FGBs on average had lower completion and higher contamination than the SGBs, all MAGs met the MIMAG standard for medium quality genomes (40) and the FGBs actually had a higher completeness and lower contigs/Mbp than GGBs (Figure 5B-E).

**Fig 5.**
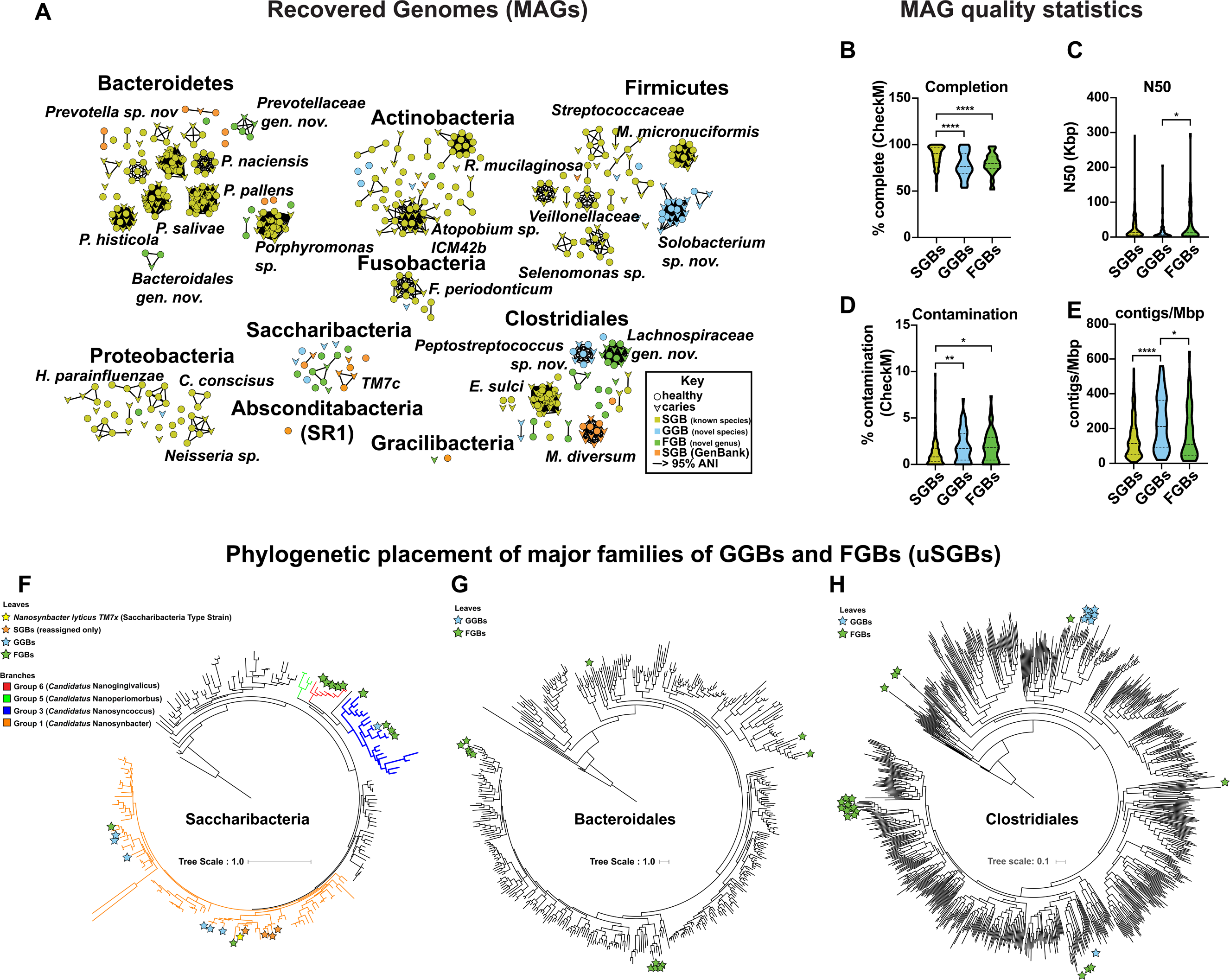
527 metagenome-assembled genomes (MAGs) were recovered. **(A) Recovery of 151 species-level genome bins (SGBs), representing 527 MAGs and ∼50 novel taxa.** Network representing an average nucleotide identity (ANI) distance matrix, generated by fastANI (42). Nodes represent MAGs, while edges represent an ANI > 95% (the cutoff chosen to designate species boundaries in this study). Circular nodes indicate MAGs recovered from healthy samples, while chevrons indicate MAGs recovered from caries samples. Nodes are colored based upon bin designation: species-level (SGB: known species; yellow), genus-level (GGB: known genus, novel species; blue), family-level (FGB: novel genus and species, green), or reassigned to SGB (described in Fig. 1; orange, Table S). Sub-networks of interest are labeled with taxonomic names. **(B-E) Statistics indicating MAG quality.** Violin charts illustrating the Completion **(B)**, N50 **(C)**, Contamination **(D)**, and contigs/Mbp **(E)** of the SGBs, GGBs, and FGBs. (completion and contamination were determined by CheckM (90)). Asterisks indicate statistical significance between indicated groups, based upon a Tukey’s Multiple Comparisons Test following a one-way ANOVA. *, p < 0.05; **, p < 0.01; ****, p < 0.0001 **(F-H) Phylogenetic trees of Saccharibacteria (F), Bacteroidales (G), and Clostridiales (H) reference genomes with placement of uSGBs.** GGBs, FGBs, and SGBs (Saccharibacteria only) are denoted by stars of the indicated color. Trees were constructed using PhyloPhAn2.

PhyloPhlAn2 (41) was used to phylogenetically place the uSGBs amongst reference strains at the order or class level. 25 of the MAGs, including 6 GGB and 11 FGB, appeared to be Candidate Phyla Radiation (CPR) bacteria. This recently described supergroup is predicted to contain >35 phyla representing >15% of the diversity of all bacteria (44). CPR taxa have long been considered microbial “dark matter” and only one species has been cultivated thus far (45). CPR have reduced genomes and are thought to be obligate epibionts (46). In this MAGs dataset, 22 CPR MAGs were Saccharibacteria (formerly known as TM7), while 2 CPR MAGs were Gracilibacteria and one was an Absconditabacteria (formerly known as SR1). A comprehensive phylogeny of currently available Saccharibacteria was recently reported, and novel named taxonomic hierarchies proposed (47). In the present study, the Saccharibacteria MAGs represent Groups 1 (order Nanosynbacteriaceae), 3 (order Nanosynsoccalia), and 6 (order Nanoperiomorables). A table with information regarding the CPR MAGs reported here is provided in Table S9 and the phylogenetic trees are provided in Figure 4F (for Saccharibacteria) and Figure S5.

GGBs other than the CPR included novel species within the genera *Peptostreptococcus, Solobacterium, Streptococcus, Lachnospiraceae, Campylobacter, Atopobium, Fusobacterium, Thermononospora, Schaalia, Parvimonas, Riemerella,* and *Granulicatella.* The *Peptostreptococcus* and two *Solobaceterium* GGBs were particularly interesting as they contained 8, 22, and 3 MAGs, respectively, indicating that these novel species may be somewhat widespread in the study population. FGBs represented novel genera and species within the taxonomic groups Atopobiaceae, Bacteroidales, Campylobacteriaceae, Clostridiales, Lachnospiraceae, Porphyromonadaceae, and Prevotellaceae. Details regarding the GGBs and FGBs are provided in Tables S10 and S11, respectively. Many of the non-CPR uSGBs were found in the clades Bacteroidales and Clostricidiales, and phylogenetic tress illustrating uSGB placement within these groups is provided in Figures 5G, 5H, S6, and S7.

The final set of MAGs was uploaded to the PATRIC database for annotation and curation using the PATRIC CLI (48), and are publicly available in the PATRIC database and RefSeq. (will be made live/public following acceptance for publication) iRep (49) was used to calculate the replication rates of MAGs, but no difference in replication rates was detected between caries and health (Figure S9, Supplemental Note 2)

## DISCUSSION

Dental caries has been known to be of bacterial origin for many decades. However, due to the “Great Plate Count Anomaly”, the true complexity of the oral microbiota (and that of indeed every microbiota!) has only began to be realized following the relatively recent development of culture-independent detection methods, such as second-generation sequencing technologies (5, 9, 16, 50). Of these techniques, 16S sequencing has been the most widely utilized technique in characterizing microbial communities, including those of the oral cavity associated with dental caries. However, 16S sequencing provides relatively low-resolution data that is biased, due to the PCR amplification step, and overlooks crucial information regarding the true diversity and functional capabilities of the communities present (14–16, 51). Sequencing is also used to quantify the resident taxa in microbial communities. However, sequencing provides only compositional data, which must be handled carefully to avoid generating spurious conclusions— a fact that is frequently swept under the rug by microbiome studies (10–13). Meanwhile, several recent landmark metagenomics studies, focused on the gut microbiome, have provided excellent examples of analyzing metagenomic (i.e. shotgun, whole genome sequencing) datasets, assembling quality MAGs, and have discovered a vast diversity of novel taxa (41, 52, 53). Major goals of this study were to perform relatively deep metagenomic sequencing to observe the microbiome present in the saliva of healthy children compared to the saliva of children with multiple dentin caries, and possibly identify novel taxa. The choice to sample saliva was mainly due to the ease of collection and ability to obtain sufficient sample volume for analysis (particularly for the case of the host markers). While the various microenviroments of the oral cavity have highly distinct microbial residents, and an ideal sampling scenario would examine diversity on this scale, saliva bathes all oral tissues and is generally thought to represent the overall oral composition (7). Differing sampling methods are likely to account for a sizable portion of the variability seen across oral microbiome studies regarding dental caries (7). Another goal of this study was to examine the concentration of host immunological markers present in saliva in caries versus health, and identify microbe-metabolite co-occurrences using a recently developed machine-learning-based tool to examine such relationships in compositional data (https://github.com/biocore/mmvec). This cross-talk between the oral microbiome and host molecules in dental caries, compared to health, is not well characterized.

This study examined the saliva of 47 children, 4-11 years old. Twenty-four children had good dental health, while 23 children had at least two carious lesions that had penetrated the enamel into the underlying dentin (*≥* 2mm deep dentin lesions), representing a relatively advanced disease state (1). Beta diversity of species-level taxonomy was significantly different between the caries and healthy groups. Notably, the canonical cariogenic species, *S. mutans,* was significantly associated with caries (3^rd^ most correlated species-level taxa according to supervised methods), but was found in relatively low abundances, and in only 11 of the 47 subjects. This indicates that *S. mutans,* when present, has a large influence on the pathogenicity of the oral microbiome due to its prodigious capacity to generate insoluble glucans and resultant biofilms (9, 26). Other oral microbiome studies have found *S. mutans* at low abundances (20, 21, 54), and the use of saliva in this study may explain its rarity, and possibly underestimation, here—as an exceptional biofilm-former, it is less likely to be shed from its dental plaque residence into the salivary milieu (55).

Both unsupervised and supervised methods showed that *Rothia, Neisseria* and *Haemophilus* spp. were associated with health, and several abundant *Prevotella* spp. (*P. histicola, pallens,* and *P. salivae*) were associated with disease. Although *Prevotella* spp. were elevated in disease, they were highly abundant in all the samples, and this correlation was not as dramatic as that of *Rothia* and *Haemophilus* with health, indicating that the positive effects of *Rothia* and *Haemophilus* may be more important than the negative effects of *Prevotella.* In a recent study, *Rothia dentocariosa* and *Rothia aeria* were associated with good dental health (31), while in another study *R. dentocariosa* was associated caries (56), indicating that the role of this genera in caries development remains nebulous. The genera *Rothia, Neisseria,* and *Haemophilus* were recently documented to be important mediators of cell-cell interactions within the early biofilm derived from healthy individuals (57), are among the first colonizers of the oral cavity after birth (58), and they were indeed largely health-associated in this study. As with *Rothia, Prevotella* spp. have been associated with both health and dental caries, depending on the study. Our findings are in line with several studies that associated *Prevotella* with dental caries (21, 59–61), including one that found it was the best predictor of childhood caries (59). These reports were in contrast to a study where *Prevotella* were enriched in the healthy cohort (62). The significant elevation in the ratio of *Prevotella* to *Rothia* observed here may represent a useful novel biomarker for caries, but wider studies are needed because the present study group was rather homogenous in terms of host ethnicity and geography.

Although the alpha diversity of the caries group was not significantly lower when species-level taxonomy was examined, alpha diversity was significantly reduced in the caries group when functional pathways were examined. Overall, the functional analyses indicated that the presence of several pathways is enriched in health, while there were no pathways detected by these methods that were unequivocally enriched in disease. Contributional diversity (i.e. how many taxa contribute a particular pathway to the community)(34) was reduced in the caries group. This was particularly evident in the aerobic respiration performed by the health-associated *Neisseria,* and several pathways that have been previously associated with the forestalling of caries pathogenesis including arginine (35) and BCAA pathways (37), which both release alkaline molecules that serve to buffer the environment and prevent enamel demineralization (63).

A number of previous studies have illustrated that penetration of the dental plaque infection into the dentin is associated with elevation of a number of cytokines and host signaling molecules (64–70). Several of these reports were supported here, where 10 immunological factors were observed at significantly elevated concentrations in the caries group compared to the healthy group. These molecules have an array of functions, and are likely to themselves influence the microbiota of the oral cavity (71). Microbe-host immunological marker co-occurrences have been characterized in periodontitis (72–74), but have not been previously examined in dental caries. Machine learning was employed to examine these microbe-molecule co-occurrences for the first time. Interestingly, the caries associated *Prevotella histicola, Prevotella salivae,* and *Veillonella aytpica* co-occurred frequently with EGF, IL-1RA, and TGFa, which were all themselves caries associated. While this co-occurrence data provides an obvious chicken or egg dilemma (and it is likely that this cross-talk is bi-directional), it also provides an atlas of microbe-host metabolite interactions that are most likely to be critical to the dysbiosis involved in caries pathogenesis, and which deserve more in-depth analysis. EGF was one of the markers most significantly elevated in the caries group, and has been previously documented to be incorporated into dentin and released on orthodontic force (75).

A further advantage of metagenomic sequencing is the ability to assemble MAGs, which allow further analysis of pan-genomics, and the identification of novel taxa. Excitingly, of the 527 MAGs reported in this study, 20% (98 MAGs) represented novel taxa, including 23 putative novel genera. 8 FGBs, representing new genera, were CPR bacteria, including Saccharibacteria and Gracilibacteria. Novel genera and species were also identified within more well-characterized genera including Prevotellaceae, Porphyromonadaceae. Several of these novel genomes, including uSGBs of *Peptostreptococcus, Solobacterium,* and *Lachnospiraceae* were assembled and binned from a large number of subjects independently, indicating that these unknown taxa may be widespread in the study population. The wealth of detailed genomic information provided by this study invites deeper analysis into the pan-genomes of the various SGBs, and investigation into the relationship of certain genotypes with disease status.

The execution of the genomics portion of this study highlighted several issues facing microbial genomics studies. One predicament is the large and growing number of databases containing genome sequences from which to choose from, as well as the quality of the contents of these repositories and maintenance of up-to-date naming and taxonomic information. An example was that the *Alloprevotella tannerae* genomes in this study initially annotated as *Prevotella tannerae* due to the naming used by the RefSeq database, despite the recognition of *Alloprevotella* as a distinct genus for several years (76). This is a particularly difficult issue to address, as changes to established phylogeny are frequent and occasionally controversial; in fact a recent study proposed a large overhaul of the bacterial tree of life, with 58% of taxa being reclassified (77). Timely implementation of improved phylogeny will help solve another issue noted in this study, the polyphyly of many taxonomic groups, particularly the class Clostridiales. Further, as mentioned above, the use of 95% ANI as the cutoff to define a species remains somewhat controversial, despite increased use and supporting evidence (42). There were several rare occasions in this dataset where SGBs, as defined by the 95% ANI distance matrix, included MAGs that best matched different (although closely related) RefSeq references. Whether this indicates that 95% ANI is not stringent enough (e.g. these should in fact be classified as multiple species) or too stringent (e.g. they should all be classified as the same species) is a debate beyond the scope of this work. It is also likely that different taxa have disparate pangenomic plasticities. Additionally, similar cutoffs and definitions for genus, family, etc. are even less well-established (77), leaving a large amount of room for interpretation with large scale studies where high numbers of novel taxa are described.

Overall, this study provided a plethora of data regarding the oral microbiome during dental caries, and its co-occurrences with host immunological markers. The tools utilized to analyze correlation to between taxa or functional pathways and disease status, as well as host markers, were designed specifically to be robust for the compositional data provided by sequencing. The authors envision the bioinformatics pipelines employed here are a useful template to guide further studies of the oral microbiome. Application of these analyses to larger and more diverse samples will dramatically improve our understanding of oral microbial ecology, and influence of the human host during dental caries.

## MATERIALS AND METHODS

### Ethics statement

Child participants and parents understood the nature of the study, and parents/guardians provided informed consent prior to the commencement of the study. The Ethics Committees of the School of Dentistry, University of California, Los Angeles, CA, USA and the J. Craig Venter Institute, La Jolla, CA, USA, approved the study design as well as the procedure for obtaining informed consent (IRB reference numbers: 13-001075 and 2016-226). All experiments were performed in accordance with the approved guidelines.

### Study Design

Subjects were included in the study if the subject was 3 years old or older, in good general health according to a medical history and clinical judgment of the clinical investigator, and had at least 12 teeth. Subjects were excluded from the study if they had generalized rampant dental caries, chronic systemic disease, or medical conditions that would influence the ability to participate in the proposed study (i.e., cancer treatment, HIV, rheumatic conditions, history of oral candidiasis). Subjects were also excluded it they had open sores or ulceration in the mouth, radiation therapy to the head and neck region of the body, significantly reduced saliva production or had been treated by anti-inflammatory or antibiotic therapy in the past 6 months. Ethnic origin was mixed for healthy subjects (Hispanic, Asian, Caucasian, Caucasian/Asian), while children with caries were of Hispanic origin. For the latter group, no other ethnic group enrolled despite several attempts to identify interested families/participants. Children with both primary and mixed dentition stages were included (caries group: 18 children with mixed dentition and 6 with primary dentition; healthy group: 19 children with mixed dentition and 6 with primary dentition). To further enable classification of health status (here caries and healthy), a comprehensive oral examination of each subject was performed as described below. Subjects were dichotomized into two groups: caries free (dmft/DMFT = 0) and caries active (subjects with ≥2 active dentin lesions). If the subject qualified for the study, (s)he was to abstain from oral hygiene activity, and eating and drinking for 2 hours prior to saliva collection in the morning. An overview of the subjects and associated metadata is provided in Table S1.

i. *Comprehensive oral examination and study groups.* The exam was performed by a single calibrated pediatric dental resident (RA), using a standard dental mirror, illuminated by artificial light. The visual inspection was aided by tactile inspection with a community periodontal index (CPI) probe when necessary. Radiographs (bitewings) were taken to determine the depth of carious lesions. The number of teeth present was recorded and their dental caries status was recorded using decayed (d), missing due to decay (m), or filled (f) teeth in primary and permanent dentitions (dmft/DMFT), according to the criteria proposed by the World Health Organization (1997) (78). Duplicate examinations were performed on 5 randomly selected subjects to assess intra-examiner reliability. Subjects were dichotomized into two groups: caries free (CF; dmft/DMFT=0) and caries active (CA; subjects with ≥2 active dentin lesions). The gingival health condition of each subject was assessed using the Gingival Index (GI) (79). GI data was published previously (80). Additionally, parent/guardian of each participant completed a survey regarding oral health regimen.
ii. *Radiographic Assessment.* Bitewing radiographs were analyzed on the XDR Imaging Software (Los Angeles, CA). Lesion depth was determined with the measuring tool, and categorized as follows: E1 (radiolucency extends to outer half of enamel), E2 (radiolucency may extend to the dentinoenamel junction), D1 (radiolucency extends to the outer one-third of dentin), D2 (radiolucency extends into the middle one third of dentin), and D3 (radiolucency extends into the inner one third of dentin)(81). To calculate the depth of lesion score, the following scores were assigned to each lesion depth: E1 = 1, E2 = 2, D1 = 3, D2 = 4, and D3 = 5, afterwards a total depth score was calculated for each subject.
iii. *Saliva Collection.* Unstimulated saliva was collected between 8:00-11:00am for the salivary immunological markers analysis. Subjects were asked to abstain from oral hygiene activity, and eating and drinking for two hours prior to collection. Before collection, subjects were instructed to rinse with water to remove all saliva from the mouth. In this study, unstimulated saliva was collected for salivary immunological marker analysis, while stimulated saliva (by chewing on sterile parafilm) was collected for Illumina sequencing (to dilute and amount of human DNA and material present). 2 ml of unstimulated saliva was collected from subjects by drooling/spitting directly into a 50mL Falcon conical tube (Fisher Scientific, Pittsburg PA) at regular intervals for a period of 5-20 minutes. Saliva samples were immediately placed on ice and protease inhibitor cocktail (Sigma, MO, USA) was added at a ratio of 100uL per 1mL of saliva to avoid protein degradation. Then saliva samples were processed by centrifugation at 6,000 x g for 5 min at 4°C, and the supernatants were transferred to cryotubes. The samples were immediately frozen in liquid nitrogen and stored at −80 °C until analysis. 2 ml of stimulated saliva was collected immediately following collection of unstimulated saliva.

### Salivary Immunological Biomarker Analysis

Frozen unstimulated saliva samples were thawed and processed through high-speed ultracentrifugation to precipitate cells and mucin for extraction of proteins. Host immunological marker profiles were determined by Multiplexed Luminex bead immunoassay (Westcoast Biosciences, San Diego, CA). A total of 38 analytes were measured, the specific immune biomarkers that were studied in saliva samples included: Epidermal Growth Factor (EGF), Fibroblast Growth Factor-2 (FGF-2), Eotaxin, Transforming Growth Factor alpha (TGF-*α*), Granulocyte Colony-Stimulating Factor (G-CSF), Granulocyte-Macrophage Colony-Stimulating Factor (GM-CSF), FMS-Like Tyrosine Kinase 3 Ligand (Flt-3L), Vascular Endothelial Growth Factor (VEGF), Fractalkine, Growth-Regulated Oncogene (GRO), Monocyte-Chemotactic Protein 3 (MCP-3), Macrophage-Derived Chemokine (MDC), Interleukin-8 (IL-8), Protein 10 (IP-10), Monocyte Chemotactic Protein-1 (MCP-1), Macrophage Inflammatory Protein-1 alpha (MIP-1*α*), Macrophage Inflammatory Protein-1 beta (MIP-1*β*), Interferon Alpha2 (IFN-*α*2), Interferon gamma (IFN-y), Interleukin-1 alpha (IL-1*α*), Interleukin-1 beta (IL-1*β*), Interleukin-1 Receptor Antagonist (IL-1RA), Interleukin-2 (IL-2), Interleukin-3 (IL-3), Interleukin-4 (IL-4), Interleukin-5 (IL-5), Interleukin-6 (IL-6), Interleukin-7 (IL-7), Interleukin-9 (IL-9), Interleukin-10 (IL-10), Interleukin-12(p40) (IL-12(p40), Interleukin-12(p70) (IL-12(p70)), Interleukin-13 (IL-13), Interleukin-15 (IL-15), Interleukin-17 (IL-17), Soluble CD40 Ligand (sCD40L), Tumor Necrosis Factor-alpha (TNF-*α*), Tumor Necrosis Factor-beta (TNF-*β*). Quantities of each host marker were compared between healthy and caries groups. In the cases of eotaxin, sCD40L, IL-17A, IL-9, IL-2, IL-3, and IL-4, the majority of samples contained levels of the respective molecule below the limit of detection for the assay. Therefore, these salivary immunological markers were not analyzed subsequently. After removal of outliers using the ROUT method with a Q = 1%, a Welch’s t-test was used to determine significantly differentially abundant immunological markers.

### DNA Extraction and sequencing

Frozen stimulated saliva samples were thawed on ice. DNA was extracted and purified from the supernatant by employing QIAmp microbiome (Qiagen) and DNA clean & concentrator (Zymo Research) kit procedures where host nucleic acid depletion step was skipped to maximize bacterial DNA recovery. Libraries were prepared using Illumina NexteraXT DNA library preparation kit according to the manufacturer’s instructions. Sequencing was carried out at the J. Craig Venter Institute (JCVI) Joint Technology Center (JTC) by using an Illumina NextSeq 500 platform (San Diego, CA, USA) (150 bp paired end reads). DNA sample concentrations were normalized at prior to sequencing. For 45 of the 47 samples, sequencing depth was 5-31 million reads per sample. Two samples, SC40 (caries) and SC33 (healthy) were sequenced ultra-deep, to 366 and 390 million reads, respectively, to examine the what information can be gleaned from even deeper sequencing. The number of reads is listed in Table S12.

### Bioinformatics analysis

i. *Quality Control.* Raw Illumina reads were subjected to quality filtering and barcode trimming using KneadData v0.5.4 (available at https://bitbucket.org/biobakery/kneaddata) by employing trimmomatic settings of 4-base wide sliding window, with average quality per base >20 and minimum length 90 bp. Reads mapping to the human genome were also removed. KneadData quality control information is provided in Table S12.
ii. *Taxonomy of reads.* Filtered reads were then analyzed using MetaPhlAn2 v2.7.5 (28) to determine relative abundances of taxa. A custom script was used to obtain an estimated number of reads using the relative abundances of each taxa and the predicted total number of reads from each sample based on MetaPhlAn2.
iii. *Calculation of beta diversity with feature loadings.* The taxonomic abundance table (i.e. OTU table) generated from MetaPhlAn2 was used as input for the QIIME2 (29) plugin, DEICODE (30), which used Robust Aitchison PCA to calculate beta diversity with feature loadings. The resulting biplot was visualized using the QIIME2 plugin Emperor (39). The feature loadings for Axis 2 of the biplot (the axis with the most difference in disease status) were visualized using Qurro (doi:10.5281/zenodo.3369454).
iv. *Using reference frames to identify taxa associated with disease.* The OTU table generated by MetaPhlAn2 was used as input for Songbird (13) in order to rank species association with disease status. The following parameters were used: number of random test examples: 5, epochs: 50,000, batch size: 3, differential-prior: 1, learning rate: 0.001. The resulting differentials were visualized using Qurro.
v. *Functional profiling of metagenomes.* HUMAnN2 (34) was used to provide information about the functional pathways present in the metagenomes. DEICODE was utilized analyze the relationship between particular metabolic pathways and disease status and Emperor was used to visualize the resulting ordination. Songbird was utilized to rank the pathways in terms of association with disease status using the same parameters described above and the ranks were visualized with Qurro.
vi. *Estimation of species-saliva immunological marker co-occurrence*. The co-occurrence of species and immunological markers was estimated using neural networks via mmvec (https://github.com/biocore/mmvec). The following parameters were used: number of testing examples: 5, minimum feature count: 10, epochs: 1000, batch size: 3, latent dim: 3, input prior: 1, output prior: 1, learning rate 0.001. Emperor was used to visualize the resulting biplot.
vii. *Assembly and binning of MAGs.* metaSPAdes was utilized to *de novo* assemble metagenomes from the quality-filtered Illumina reads (82). The resulting assemblies were binned using the MetaWRAP pipeline v1.1.5 (83). The MetaWRAP initial_binning module used Maxbin2, Metabat2, and Concoct. Subsequently, the bin_refinement module was used to construct the best final bin set by comparing the results of the 3 binning tools. The bin_reassembly module was then used to reassemble the final bin set to make further improvements. The quality control cutoffs for all MetaWRAP modules were >50% completeness and <10% contamination, which are the cutoffs for Medium-Quality Draft Metagenome-Assembled Genomes as set forth by the Genome Standards Consortium (40). This generated 527 metagenome-assembled genomes (MAGs) that were at least of medium quality. The classify_bins and quant_bins modules were used to respectively obtain a taxonomy estimate based on megablast and to provide the quantity of each bin in the form of ‘genome copies per million reads’.
viii. *Determining species level genome bins.* To identify species-level genome bins (SGBs), Mash v2.1 (43) was used to query all 527 MAGs against the entire RefSeq database with a Mash distance cutoff of 5 (corresponding to a 95% average nucleotide identity (ANI)). Although a topic of some debate, 95%ANI has been used by several recent landmark studies as the cutoff for genomes representing the same species. All MAGs with a RefSeq hit with a Mash distance of <5 were assigned the species name of that hit. Because Mash can underestimate ANI for less than complete genomes, fastANI v1.1 was used to compare the ANI of all 570 MAGs and generate a distance matrix. This distance matrix was used to create a Cytoscape network to visualize all MAGs that had an ANI>95% (i.e. link all bins that were of the same species, based on ANI, with an edge). The fact that Mash distance <5 and fastANI ANI>95% aligned almost perfectly served as a useful internal control. This strategy resulted in 151 SGB’s (91 with no connection and 60 with at least one). 95 SGBs, representing 444 total bins (MAGs), had a Mash and/or fastANI hit with ANI >95%, and were termed known SGBs (kSGBs). 56 of the total SGBs, representing 126 total bins (MAGs), did not have hit in RefSeq with at least 95%ANI and therefore were defined as unknown SGBs (uSGBs). To further define the uSGBs, a combination of Mash distances < 30 to the RefSeq database, CheckM, the metaWRAP classify_bins module (which uses taxator TK), fastANI, Kraken, and blastx were used to determine the taxonomy of the uSGBs. When uSGBs contained multiple MAGs, the MAG with the best quality score according to the formula (completion – (2x contamination)) was used to find the best hit. Previous studies have utilized ANI cutoffs of 85% and 70% to determine genus and family level genome bins, and a similar approach was used here with manual evaluation of the closest ANI hits leading to assignment of uSGBs to genus-level genome bins (GGBs) or family level genome bins (FGBs).
ix. *Phylogenetic placement of uSGBs.* PhyloPhlAn2 (41) was used to determine the phylogeny of uSGBs. The following parameters were used: --diversity medium –accurate. The following external tools were used: diamond (84), mafft (85), trimal (86), fasttree (87), and RAxML (88). Resulting phylogenetic trees were visualized using iToL 4.(89)
x. *Inference of actively replicating taxa*. iRep (49) was utilized to infer the replication rates of taxa in the metagenomes assembled using metaSPAdes as described above. iRep was then used to calculate the estimated replication rate of genera for which sufficient draft genomes had been assembled from the metagenomic data.

## Data availability

Sequence data have been submitted to NCBI under BioProject ID PRJNA478018 with SRA accession number SRP151559. MAGs have been uploaded to PATRIC and annotated (will be made public upon acceptance for publication).

## ACKNOWLEDGEMENTS

This study was supported by NIH/NIDCR grants F32-DE026947 (J.L.B.) and R00-DE024543 (A.E.). The authors also acknowledge and thank R. Alexander Richter, Semar Petrus, Josh Espinoza, Drishti Kaul, Clarisse Marotz, Marcus Fedarko, and Cameron Martino for very helpful scientific discussions.

## REFERENCES

1. Pitts NB, Zero DT, Marsh PD, Ekstrand K, Weintraub JA, Ramos-Gomez F, Tagami J, Twetman S, Tsakos G, Ismail A. 2017. Dental caries. Nat Rev Dis Primers 3:17030.

2. Loesche WJ. 1986. Role of Streptococcus mutans in human dental decay. Microbiol Rev 50:353–80.

3. Simon-Soro A, Mira A. 2015. Solving the etiology of dental caries. Trends Microbiol 23:76–82.

4. Bowen WH, Burne RA, Wu H, Koo H. 2018. Oral Biofilms: Pathogens, Matrix, and Polymicrobial Interactions in Microenvironments. Trends Microbiol 26:229–242.

5. Burne RA. 2018. Getting to Know “The Known Unknowns”: Heterogeneity in the Oral Microbiome. Adv Dent Res 29:66–70.

6. Cross B, Faustoferri RC, Quivey RG, Jr. 2016. What are We Learning and What Can We Learn from the Human Oral Microbiome Project? Curr Oral Health Rep 3:56–63.

7. Mira A. 2018. Oral Microbiome Studies: Potential Diagnostic and Therapeutic Implications. Adv Dent Res 29:71–77.

8. Philip N, Suneja B, Walsh L. 2018. Beyond Streptococcus mutans: clinical implications of the evolving dental caries aetiological paradigms and its associated microbiome. Br Dent J 224:219–225.

9. Banas JA, Drake DR. 2018. Are the mutans streptococci still considered relevant to understanding the microbial etiology of dental caries? BMC Oral Health 18:129.

10. Gloor GB, Macklaim JM, Pawlowsky-Glahn V, Egozcue JJ. 2017. Microbiome Datasets Are Compositional: And This Is Not Optional. Front Microbiol 8:2224.

11. Morton JT, Sanders J, Quinn RA, McDonald D, Gonzalez A, Vazquez-Baeza Y, Navas-Molina JA, Song SJ, Metcalf JL, Hyde ER, Lladser M, Dorrestein PC, Knight R. 2017. Balance Trees Reveal Microbial Niche Differentiation. mSystems 2.

12. Knight R, Vrbanac A, Taylor BC, Aksenov A, Callewaert C, Debelius J, Gonzalez A, Kosciolek T, McCall LI, McDonald D, Melnik AV, Morton JT, Navas J, Quinn RA, Sanders JG, Swafford AD, Thompson LR, Tripathi A, Xu ZZ, Zaneveld JR, Zhu Q, Caporaso JG, Dorrestein PC. 2018. Best practices for analysing microbiomes. Nat Rev Microbiol 16:410–422.

13. Morton JT, Marotz C, Washburne A, Silverman J, Zaramela LS, Edlund A, Zengler K, Knight R. 2019. Establishing microbial composition measurement standards with reference frames. Nat Commun 10:2719.

14. Hong S, Bunge J, Leslin C, Jeon S, Epstein SS. 2009. Polymerase chain reaction primers miss half of rRNA microbial diversity. ISME J 3:1365–73.

15. Pinto AJ, Raskin L. 2012. PCR biases distort bacterial and archaeal community structure in pyrosequencing datasets. PLoS One 7:e43093.

16. Jovel J, Patterson J, Wang W, Hotte N, O’Keefe S, Mitchel T, Perry T, Kao D, Mason AL, Madsen KL, Wong GK. 2016. Characterization of the Gut Microbiome Using 16S or Shotgun Metagenomics. Front Microbiol 7:459.

17. Louca S, Doebeli M, Parfrey LW. 2018. Correcting for 16S rRNA gene copy numbers in microbiome surveys remains an unsolved problem. Microbiome 6:41.

18. Human Microbiome Project C. 2012. Structure, function and diversity of the healthy human microbiome. Nature 486:207–14.

19. Langille MG, Zaneveld J, Caporaso JG, McDonald D, Knights D, Reyes JA, Clemente JC, Burkepile DE, Vega Thurber RL, Knight R, Beiko RG, Huttenhower C. 2013. Predictive functional profiling of microbial communities using 16S rRNA marker gene sequences. Nat Biotechnol 31:814–21.

20. Espinoza JL, Harkins DM, Torralba M, Gomez A, Highlander SK, Jones MB, Leong P, Saffery R, Bockmann M, Kuelbs C, Inman JM, Hughes T, Craig JM, Nelson KE, Dupont CL. 2018. Supragingival Plaque Microbiome Ecology and Functional Potential in the Context of Health and Disease. MBio 9.

21. Al-Hebshi NN, Baraniya D, Chen T, Hill J, Puri S, Tellez M, Hasan NA, Colwell RR, Ismail A. 2019. Metagenome sequencing-based strain-level and functional characterization of supragingival microbiome associated with dental caries in children. J Oral Microbiol 11:1557986.

22. Belda-Ferre P, Alcaraz LD, Cabrera-Rubio R, Romero H, Simon-Soro A, Pignatelli M, Mira A. 2012. The oral metagenome in health and disease. ISME J 6:46–56.

23. Belstrom D, Constancias F, Liu Y, Yang L, Drautz-Moses DI, Schuster SC, Kohli GS, Jakobsen TH, Holmstrup P, Givskov M. 2017. Metagenomic and metatranscriptomic analysis of saliva reveals disease-associated microbiota in patients with periodontitis and dental caries. NPJ Biofilms Microbiomes 3:23.

24. Costalonga M, Herzberg MC. 2014. The oral microbiome and the immunobiology of periodontal disease and caries. Immunol Lett 162:22–38.

25. Meyle J, Dommisch H, Groeger S, Giacaman RA, Costalonga M, Herzberg M. 2017. The innate host response in caries and periodontitis. J Clin Periodontol 44:1215–1225.

26. Bowen WH. 2016. Dental caries - not just holes in teeth! A perspective. Mol Oral Microbiol 31:228–33.

27. Baker JL, Edlund A. 2018. Exploiting the Oral Microbiome to Prevent Tooth Decay: Has Evolution Already Provided the Best Tools? Front Microbiol 9:3323.

28. Truong DT, Franzosa EA, Tickle TL, Scholz M, Weingart G, Pasolli E, Tett A, Huttenhower C, Segata N. 2015. MetaPhlAn2 for enhanced metagenomic taxonomic profiling. Nat Methods 12:902–3.

29. Bolyen E, Rideout JR, Dillon MR, Bokulich NA, Abnet CC, Al-Ghalith GA, Alexander H, Alm EJ, Arumugam M, Asnicar F, Bai Y, Bisanz JE, Bittinger K, Brejnrod A, Brislawn CJ, Brown CT, Callahan BJ, Caraballo-Rodriguez AM, Chase J, Cope EK, Da Silva R, Diener C, Dorrestein PC, Douglas GM, Durall DM, Duvallet C, Edwardson CF, Ernst M, Estaki M, Fouquier J, Gauglitz JM, Gibbons SM, Gibson DL, Gonzalez A, Gorlick K, Guo J, Hillmann B, Holmes S, Holste H, Huttenhower C, Huttley GA, Janssen S, Jarmusch AK, Jiang L, Kaehler BD, Kang KB, Keefe CR, Keim P, Kelley ST, Knights D, et al. 2019. Reproducible, interactive, scalable and extensible microbiome data science using QIIME 2. Nat Biotechnol 37:852–857.

30. Martino C, Morton JT, Marotz CA, Thompson LR, Tripathi A, Knight R, Zengler K. 2019. A Novel Sparse Compositional Technique Reveals Microbial Perturbations. mSystems 4.

31. Agnello M, Marques J, Cen L, Mittermuller B, Huang A, Chaichanasakul Tran N, Shi W, He X, Schroth RJ. 2017. Microbiome Associated with Severe Caries in Canadian First Nations Children. J Dent Res 96:1378–1385.

32. Thomas RZ, Zijnge V, Cicek A, de Soet JJ, Harmsen HJ, Huysmans MC. 2012. Shifts in the microbial population in relation to in situ caries progression. Caries Res 46:427–31.

33. Pereira D, Seneviratne CJ, Koga-Ito CY, Samaranayake LP. 2018. Is the oral fungal pathogen Candida albicans a cariogen? Oral Dis 24:518–526.

34. Franzosa EA, McIver LJ, Rahnavard G, Thompson LR, Schirmer M, Weingart G, Lipson KS, Knight R, Caporaso JG, Segata N, Huttenhower C. 2018. Species-level functional profiling of metagenomes and metatranscriptomes. Nat Methods 15:962–968.

35. Nascimento MM, Alvarez AJ, Huang X, Hanway S, Perry S, Luce A, Richards VP, Burne RA. 2019. Arginine Metabolism in Supragingival Oral Biofilms as a Potential Predictor of Caries Risk. JDR Clin Trans Res 4:262–270.

36. Fozo EM, Scott-Anne K, Koo H, Quivey RG, Jr. 2007. Role of unsaturated fatty acid biosynthesis in virulence of Streptococcus mutans. Infect Immun 75:1537–9.

37. Santiago B, MacGilvray M, Faustoferri RC, Quivey RG, Jr. 2012. The branched-chain amino acid aminotransferase encoded by ilvE is involved in acid tolerance in Streptococcus mutans. J Bacteriol 194:2010–9.

38. Liu YL, Nascimento M, Burne RA. 2012. Progress toward understanding the contribution of alkali generation in dental biofilms to inhibition of dental caries. Int J Oral Sci 4:135–40.

39. Vazquez-Baeza Y, Pirrung M, Gonzalez A, Knight R. 2013. EMPeror: a tool for visualizing high-throughput microbial community data. Gigascience 2:16.

40. Bowers RM, Kyrpides NC, Stepanauskas R, Harmon-Smith M, Doud D, Reddy TBK, Schulz F, Jarett J, Rivers AR, Eloe-Fadrosh EA, Tringe SG, Ivanova NN, Copeland A, Clum A, Becraft ED, Malmstrom RR, Birren B, Podar M, Bork P, Weinstock GM, Garrity GM, Dodsworth JA, Yooseph S, Sutton G, Glockner FO, Gilbert JA, Nelson WC, Hallam SJ, Jungbluth SP, Ettema TJG, Tighe S, Konstantinidis KT, Liu WT, Baker BJ, Rattei T, Eisen JA, Hedlund B, McMahon KD, Fierer N, Knight R, Finn R, Cochrane G, Karsch-Mizrachi I, Tyson GW, Rinke C, Genome Standards C, Lapidus A, Meyer F, Yilmaz P, Parks DH, et al. 2017. Minimum information about a single amplified genome (MISAG) and a metagenome-assembled genome (MIMAG) of bacteria and archaea. Nat Biotechnol 35:725–731.

41. Pasolli E, Asnicar F, Manara S, Zolfo M, Karcher N, Armanini F, Beghini F, Manghi P, Tett A, Ghensi P, Collado MC, Rice BL, DuLong C, Morgan XC, Golden CD, Quince C, Huttenhower C, Segata N. 2019. Extensive Unexplored Human Microbiome Diversity Revealed by Over 150,000 Genomes from Metagenomes Spanning Age, Geography, and Lifestyle. Cell 176:649–662 e20.

42. Jain C, Rodriguez RL, Phillippy AM, Konstantinidis KT, Aluru S. 2018. High throughput ANI analysis of 90K prokaryotic genomes reveals clear species boundaries. Nat Commun 9:5114.

43. Ondov BD, Treangen TJ, Melsted P, Mallonee AB, Bergman NH, Koren S, Phillippy AM. 2016. Mash: fast genome and metagenome distance estimation using MinHash. Genome Biol 17:132.

44. Hug LA, Baker BJ, Anantharaman K, Brown CT, Probst AJ, Castelle CJ, Butterfield CN, Hernsdorf AW, Amano Y, Ise K, Suzuki Y, Dudek N, Relman DA, Finstad KM, Amundson R, Thomas BC, Banfield JF. 2016. A new view of the tree of life. Nat Microbiol 1:16048.

45. He X, McLean JS, Edlund A, Yooseph S, Hall AP, Liu SY, Dorrestein PC, Esquenazi E, Hunter RC, Cheng G, Nelson KE, Lux R, Shi W. 2015. Cultivation of a human-associated TM7 phylotype reveals a reduced genome and epibiotic parasitic lifestyle. Proc Natl Acad Sci U S A 112:244–9.

46. Baker JL, Bor B, Agnello M, Shi W, He X. 2017. Ecology of the Oral Microbiome: Beyond Bacteria. Trends Microbiol 25:362–374.

47. McLean JS, Bor B, To TT, Liu Q, Kerns KA, Solden L, Wrighton K, He X, Shi W. 2018. Evidence of independent acquisition and adaption of ultra-small bacteria to human hosts across the highly diverse yet reduced genomes of the phylum Saccharibacteria. bioRxiv doi:10.1101/258137:258137.

48. Wattam AR, Davis JJ, Assaf R, Boisvert S, Brettin T, Bun C, Conrad N, Dietrich EM, Disz T, Gabbard JL, Gerdes S, Henry CS, Kenyon RW, Machi D, Mao C, Nordberg EK, Olsen GJ, Murphy-Olson DE, Olson R, Overbeek R, Parrello B, Pusch GD, Shukla M, Vonstein V, Warren A, Xia F, Yoo H, Stevens RL. 2017. Improvements to PATRIC, the all-bacterial Bioinformatics Database and Analysis Resource Center. Nucleic Acids Res 45:D535–D542.

49. Brown CT, Olm MR, Thomas BC, Banfield JF. 2016. Measurement of bacterial replication rates in microbial communities. Nat Biotechnol 34:1256–1263.

50. Kilian M, Chapple IL, Hannig M, Marsh PD, Meuric V, Pedersen AM, Tonetti MS, Wade WG, Zaura E. 2016. The oral microbiome - an update for oral healthcare professionals. Br Dent J 221:657–666.

51. Hillmann B, Al-Ghalith GA, Shields-Cutler RR, Zhu Q, Gohl DM, Beckman KB, Knight R, Knights D. 2018. Evaluating the Information Content of Shallow Shotgun Metagenomics. mSystems 3.

52. Almeida A, Mitchell AL, Boland M, Forster SC, Gloor GB, Tarkowska A, Lawley TD, Finn RD. 2019. A new genomic blueprint of the human gut microbiota. Nature 568:499–504.

53. Nayfach S, Shi ZJ, Seshadri R, Pollard KS, Kyrpides NC. 2019. New insights from uncultivated genomes of the global human gut microbiome. Nature 568:505–510.

54. Simon-Soro A, Belda-Ferre P, Cabrera-Rubio R, Alcaraz LD, Mira A. 2013. A tissue-dependent hypothesis of dental caries. Caries Res 47:591–600.

55. Lemos JA, Palmer SR, Zeng L, Wen ZT, Kajfasz JK, Freires IA, Abranches J, Brady LJ. 2019. The Biology of Streptococcus mutans. Microbiol Spectr 7.

56. Jiang S, Gao X, Jin L, Lo EC. 2016. Salivary Microbiome Diversity in Caries-Free and Caries-Affected Children. Int J Mol Sci 17.

57. Palmer RJ, Jr., Shah N, Valm A, Paster B, Dewhirst F, Inui T, Cisar JO. 2017. Interbacterial Adhesion Networks within Early Oral Biofilms of Single Human Hosts. Appl Environ Microbiol 83.

58. Sulyanto RM, Thompson ZA, Beall CJ, Leys EJ, Griffen AL. 2019. The Predominant Oral Microbiota Is Acquired Early in an Organized Pattern. Sci Rep 9:10550.

59. Teng F, Yang F, Huang S, Bo C, Xu ZZ, Amir A, Knight R, Ling J, Xu J. 2015. Prediction of Early Childhood Caries via Spatial-Temporal Variations of Oral Microbiota. Cell Host Microbe 18:296–306.

60. Hurley E, Barrett MPJ, Kinirons M, Whelton H, Ryan CA, Stanton C, Harris HMB, O’Toole PW. 2019. Comparison of the salivary and dentinal microbiome of children with severe-early childhood caries to the salivary microbiome of caries-free children. BMC Oral Health 19:13.

61. Tanner AC, Kent RL, Jr., Holgerson PL, Hughes CV, Loo CY, Kanasi E, Chalmers NI, Johansson I. 2011. Microbiota of severe early childhood caries before and after therapy. J Dent Res 90:1298–305.

62. Gomez A, Espinoza JL, Harkins DM, Leong P, Saffery R, Bockmann M, Torralba M, Kuelbs C, Kodukula R, Inman J, Hughes T, Craig JM, Highlander SK, Jones MB, Dupont CL, Nelson KE. 2017. Host Genetic Control of the Oral Microbiome in Health and Disease. Cell Host Microbe 22:269–278 e3.

63. Baker JL, Faustoferri RC, Quivey RG, Jr. 2017. Acid-adaptive mechanisms of Streptococcus mutans-the more we know, the more we don’t. Mol Oral Microbiol 32:107–117.

64. Adachi T, Nakanishi T, Yumoto H, Hirao K, Takahashi K, Mukai K, Nakae H, Matsuo T. 2007. Caries-related bacteria and cytokines induce CXCL10 in dental pulp. J Dent Res 86:1217–22.

65. Artese L, Rubini C, Ferrero G, Fioroni M, Santinelli A, Piattelli A. 2002. Vascular endothelial growth factor (VEGF) expression in healthy and inflamed human dental pulps. J Endod 28:20–3.

66. Kokkas AB, Goulas A, Varsamidis K, Mirtsou V, Tziafas D. 2007. Irreversible but not reversible pulpitis is associated with up-regulation of tumour necrosis factor-alpha gene expression in human pulp. Int Endod J 40:198–203.

67. McLachlan JL, Sloan AJ, Smith AJ, Landini G, Cooper PR. 2004. S100 and cytokine expression in caries. Infect Immun 72:4102–8.

68. Hahn CL, Best AM, Tew JG. 2000. Cytokine induction by Streptococcus mutans and pulpal pathogenesis. Infect Immun 68:6785–9.

69. Sloan AJ, Perry H, Matthews JB, Smith AJ. 2000. Transforming growth factor-beta isoform expression in mature human healthy and carious molar teeth. Histochem J 32:247–52.

70. Horst OV, Horst JA, Samudrala R, Dale BA. 2011. Caries induced cytokine network in the odontoblast layer of human teeth. BMC Immunol 12:9.

71. Chang AM, Liu Q, Hajjar AM, Greer A, McLean JS, Darveau RP. 2019. Toll-like receptor 2 and toll-like receptor 4 responses regulate neutrophil infiltration into the junctional epithelium and significantly contribute to the composition of the oral microbiota. J Periodontol doi:10.1002/JPER.18-0719.

72. Zhou J, Yao Y, Jiao K, Zhang J, Zheng X, Wu F, Hu X, Li J, Yu Z, Zhang G, Jiang N, Li Z. 2017. Relationship between Gingival Crevicular Fluid Microbiota and Cytokine Profile in Periodontal Host Homeostasis. Front Microbiol 8:2144.

73. Arias-Bujanda N, Regueira-Iglesias A, Alonso-Sampedro M, Gonzalez-Peteiro MM, Mira A, Balsa-Castro C, Tomas I. 2018. Cytokine Thresholds in Gingival Crevicular Fluid with Potential Diagnosis of Chronic Periodontitis Differentiating by Smoking Status. Sci Rep 8:18003.

74. Lundmark A, Hu YOO, Huss M, Johannsen G, Andersson AF, Yucel-Lindberg T. 2019. Identification of Salivary Microbiota and Its Association With Host Inflammatory Mediators in Periodontitis. Front Cell Infect Microbiol 9:216.

75. Derringer K, Linden R. 2007. Epidermal growth factor released in human dental pulp following orthodontic force. Eur J Orthod 29:67–71.

76. Downes J, Dewhirst FE, Tanner AC, Wade WG. 2013. Description of Alloprevotella rava gen. nov., sp. nov., isolated from the human oral cavity, and reclassification of Prevotella tannerae Moore et al. 1994 as Alloprevotella tannerae gen. nov., comb. nov. Int J Syst Evol Microbiol 63:1214–8.

77. Parks DH, Chuvochina M, Waite DW, Rinke C, Skarshewski A, Chaumeil PA, Hugenholtz P. 2018. A standardized bacterial taxonomy based on genome phylogeny substantially revises the tree of life. Nat Biotechnol 36:996–1004.

78. Organization WH. 1971. Oral health surveys: basic methods. World Health Organization.

79. Loe H. 1967. The Gingival Index, the Plaque Index and the Retention Index Systems. J Periodontol 38:Suppl:610–6.

80. Aleti G, Baker JL, Tang X, Alvarez R, Dinis M, Tran NC, Melnik AV, Zhong C, Ernst M, Dorrestein PC, Edlund A. 2019. Identification of the Bacterial Biosynthetic Gene Clusters of the Oral Microbiome Illuminates the Unexplored Social Language of Bacteria during Health and Disease. MBio 10.

81. Anusavice KJ. 2005. Present and future approaches for the control of caries. J Dent Educ 69:538–54.

82. Nurk S, Meleshko D, Korobeynikov A, Pevzner PA. 2017. metaSPAdes: a new versatile metagenomic assembler. Genome Res 27:824–834.

83. Uritskiy GV, DiRuggiero J, Taylor J. 2018. MetaWRAP-a flexible pipeline for genome-resolved metagenomic data analysis. Microbiome 6:158.

84. Buchfink B, Xie C, Huson DH. 2015. Fast and sensitive protein alignment using DIAMOND. Nat Methods 12:59–60.

85. Katoh K, Standley DM. 2013. MAFFT multiple sequence alignment software version 7: improvements in performance and usability. Mol Biol Evol 30:772–80.

86. Capella-Gutierrez S, Silla-Martinez JM, Gabaldon T. 2009. trimAl: a tool for automated alignment trimming in large-scale phylogenetic analyses. Bioinformatics 25:1972–3.

87. Price MN, Dehal PS, Arkin AP. 2009. FastTree: computing large minimum evolution trees with profiles instead of a distance matrix. Mol Biol Evol 26:1641–50.

88. Stamatakis A. 2014. RAxML version 8: a tool for phylogenetic analysis and post-analysis of large phylogenies. Bioinformatics 30:1312–3.

89. Letunic I, Bork P. 2019. Interactive Tree Of Life (iTOL) v4: recent updates and new developments. Nucleic Acids Res 47:W256–W259.

90. Parks DH, Imelfort M, Skennerton CT, Hugenholtz P, Tyson GW. 2015. CheckM: assessing the quality of microbial genomes recovered from isolates, single cells, and metagenomes. Genome Res 25:1043–55.

