## Supplementary figures and images for "Deep metagenomics examines the oral microbiome during dental caries, revealing novel taxa and co-occurrences with host molecules"

### Figure S1

**Figure S1**

##### Taxonomic abundance heatmap

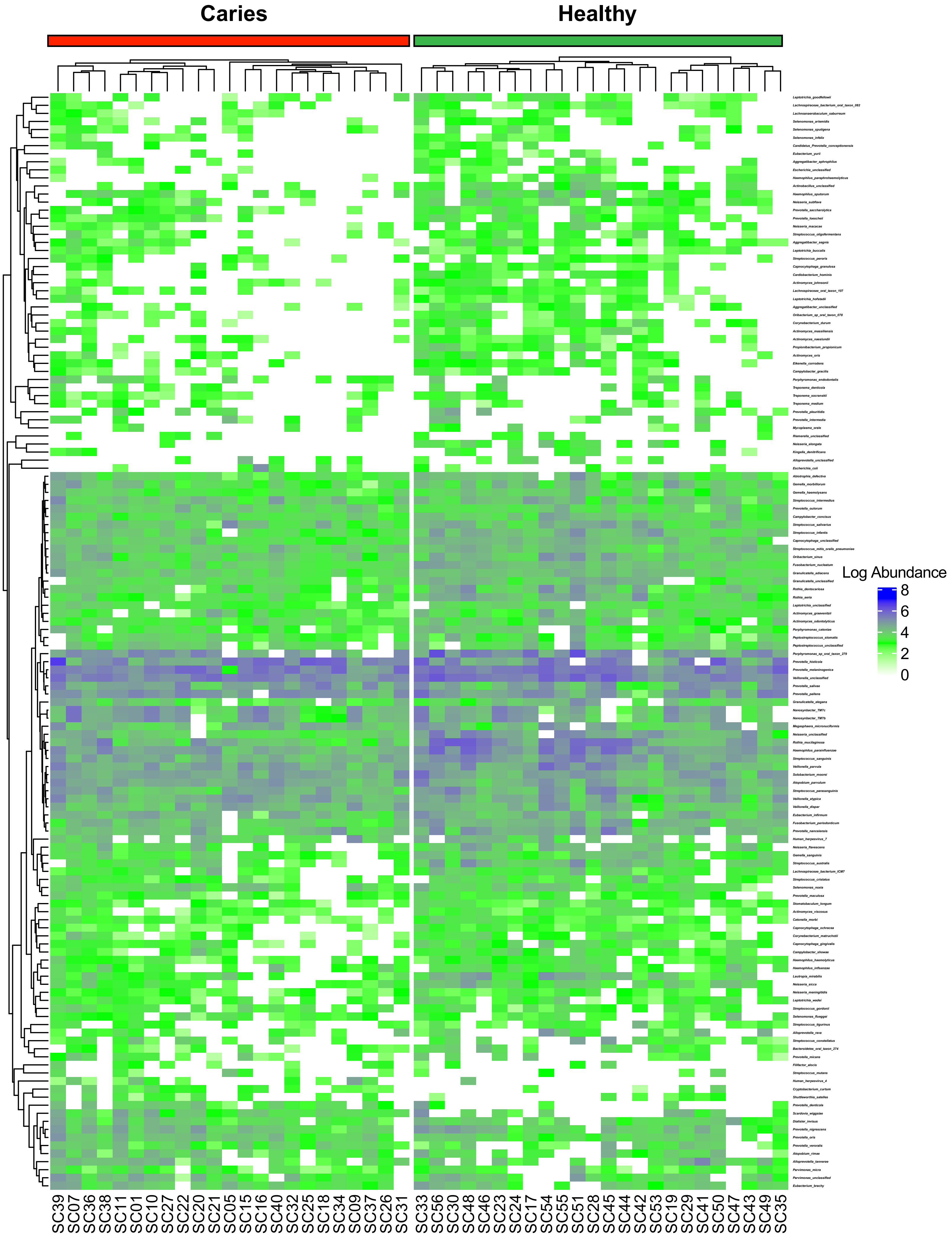

### Figure S2

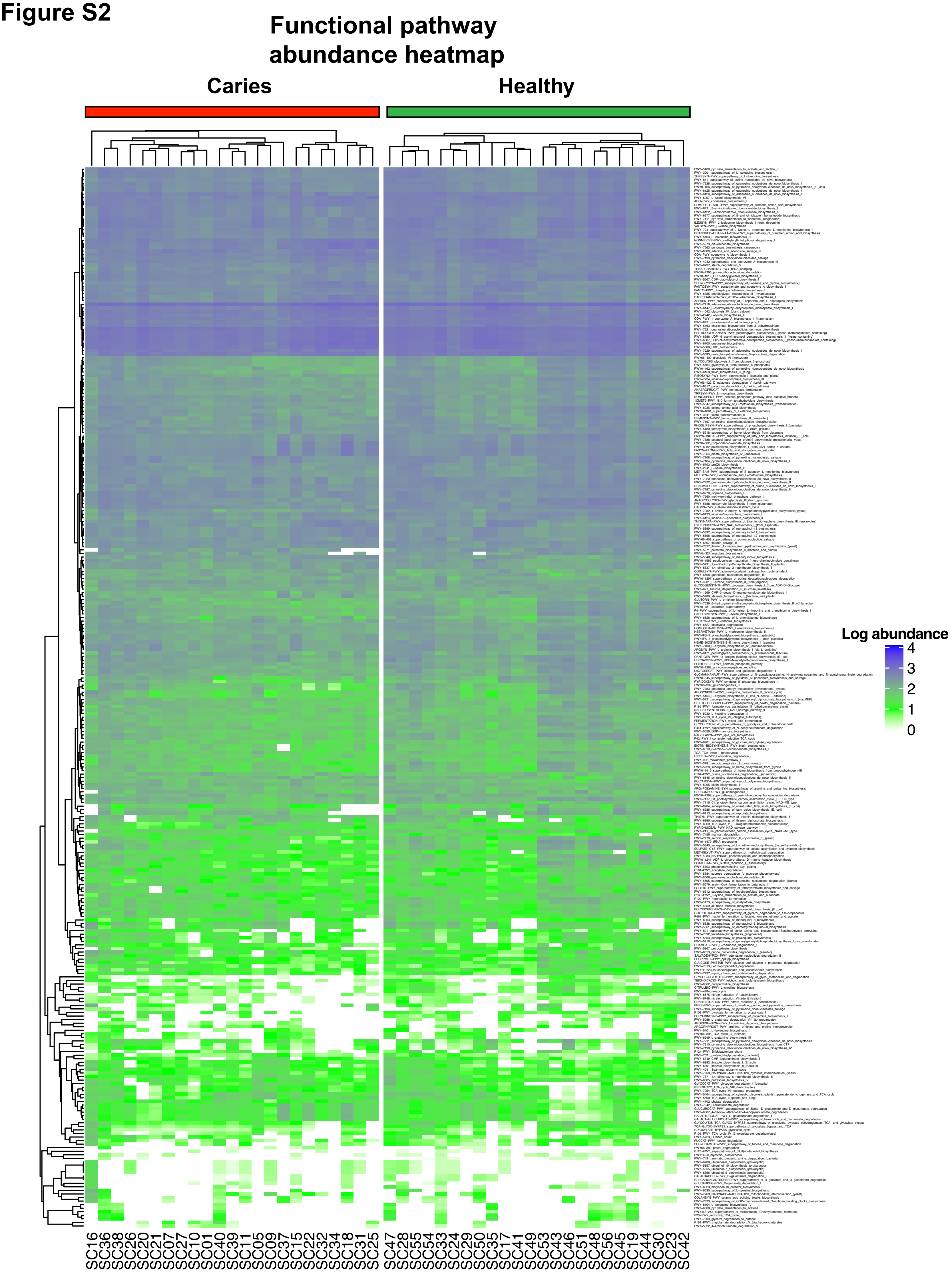

### Figure S3

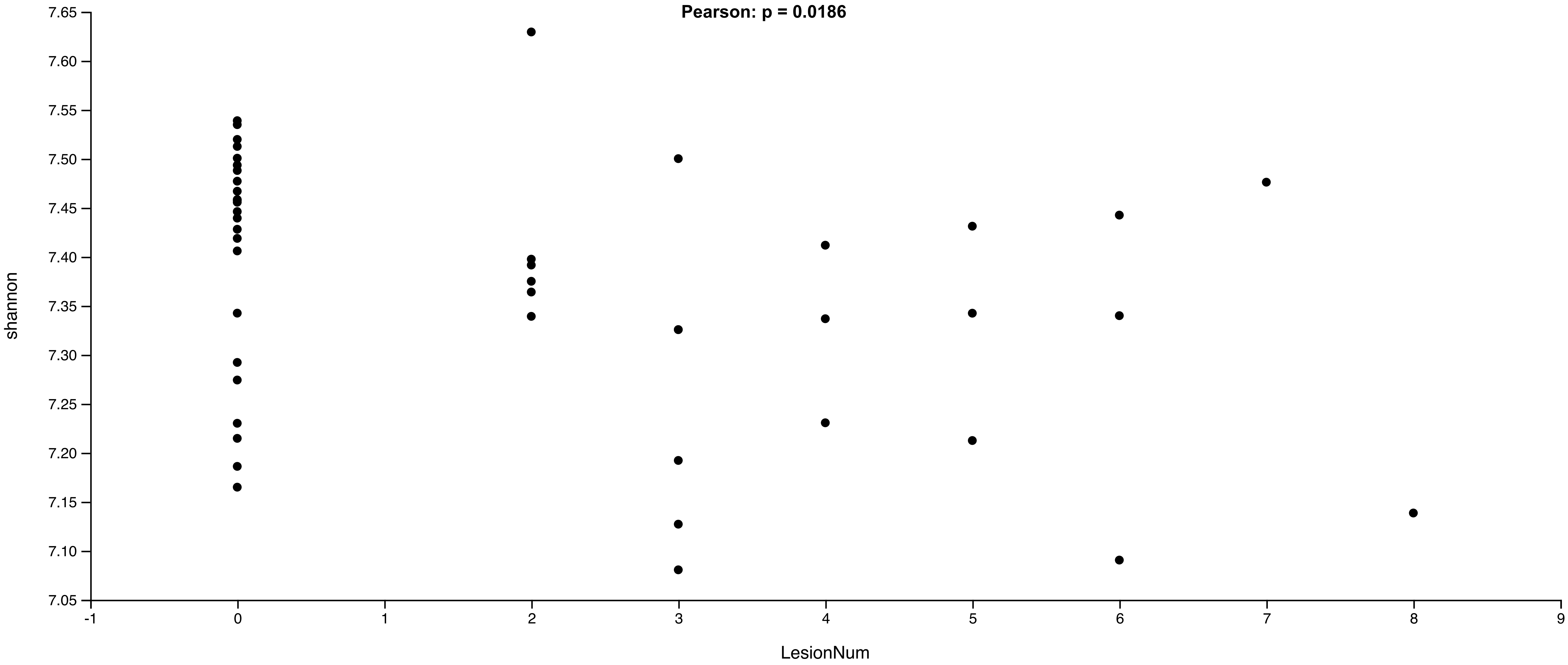

### Figure S4

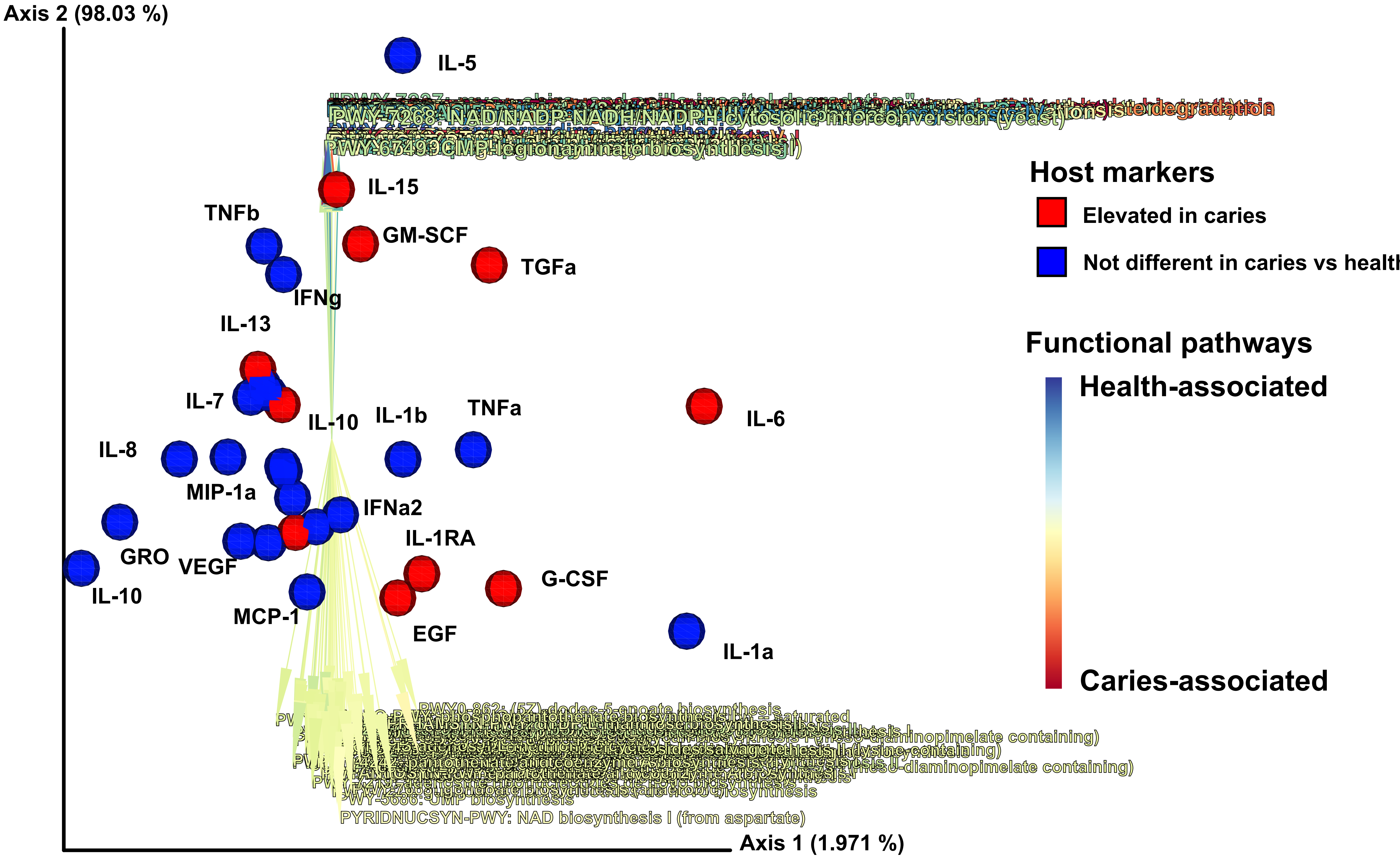

### Figure S5

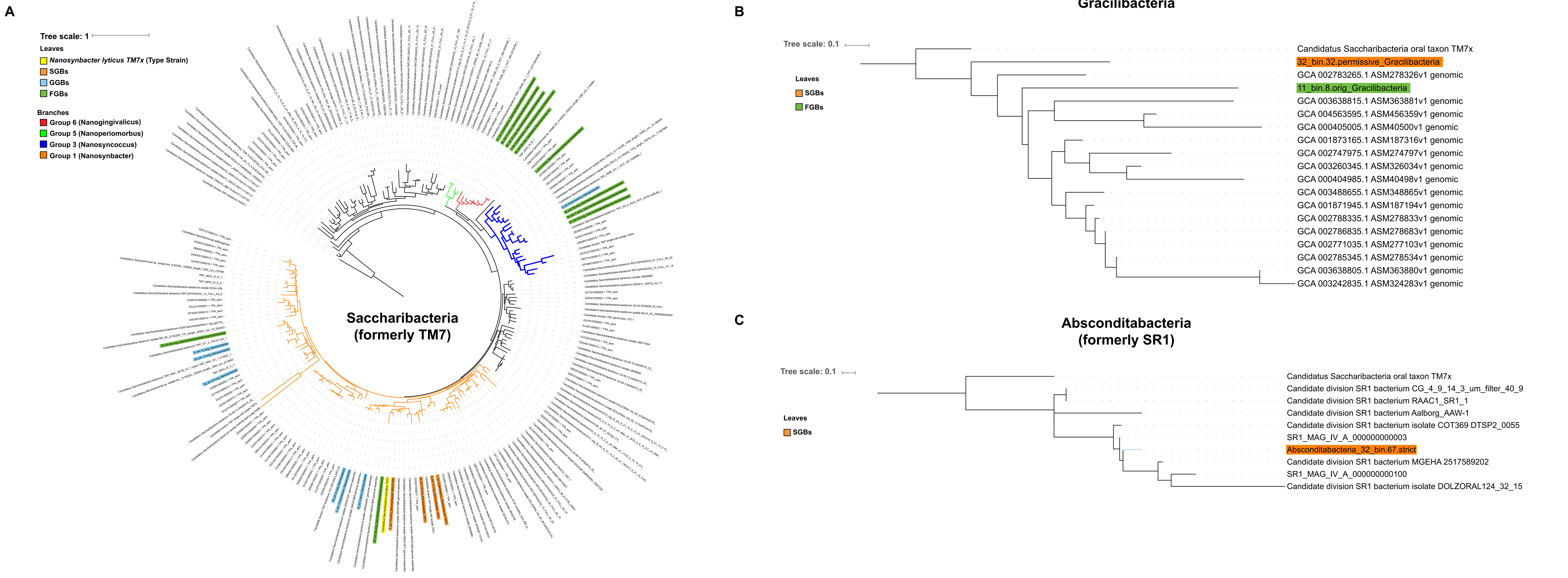

### Figure S6

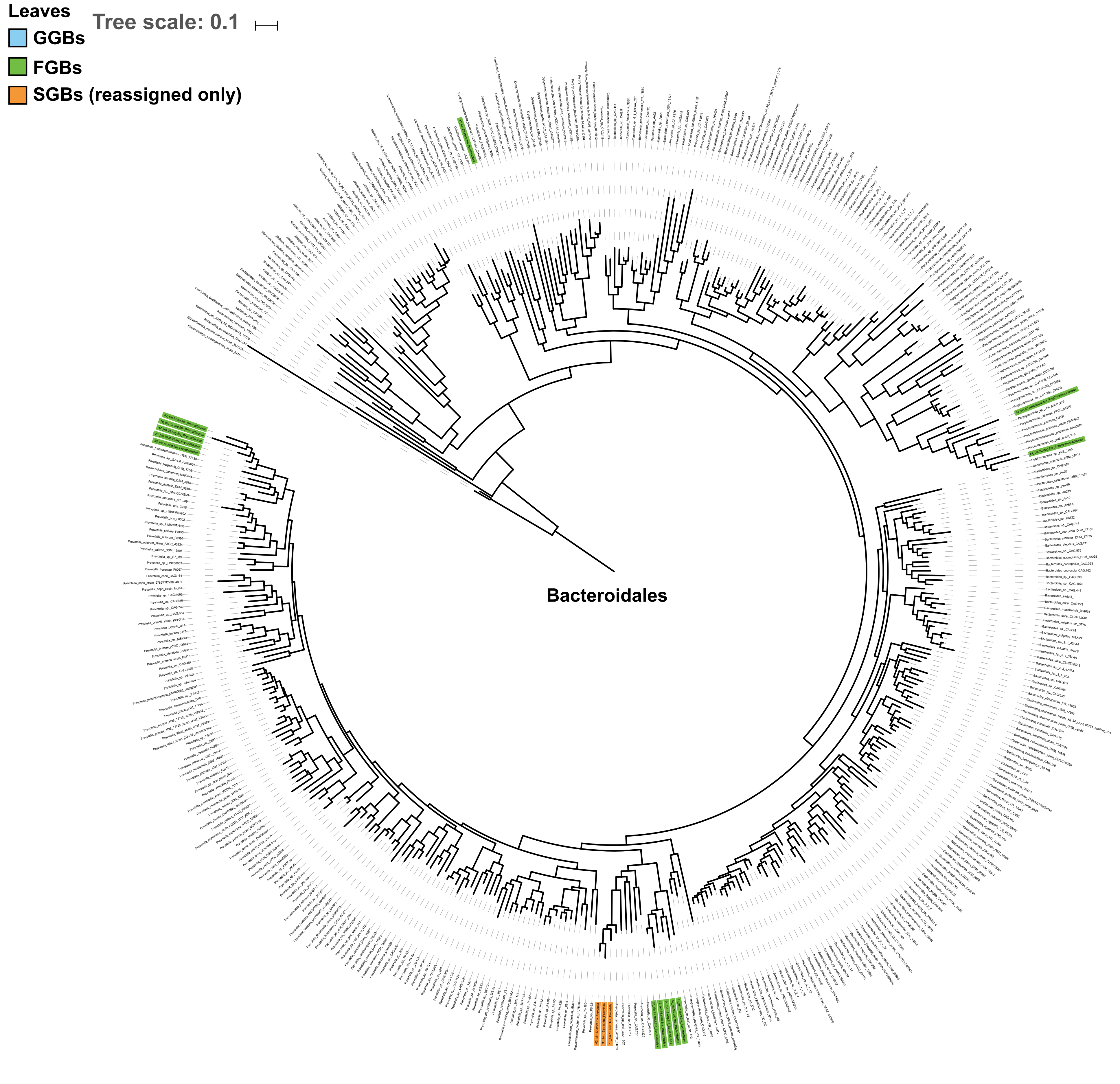

### Figure S7

#### Figure S7

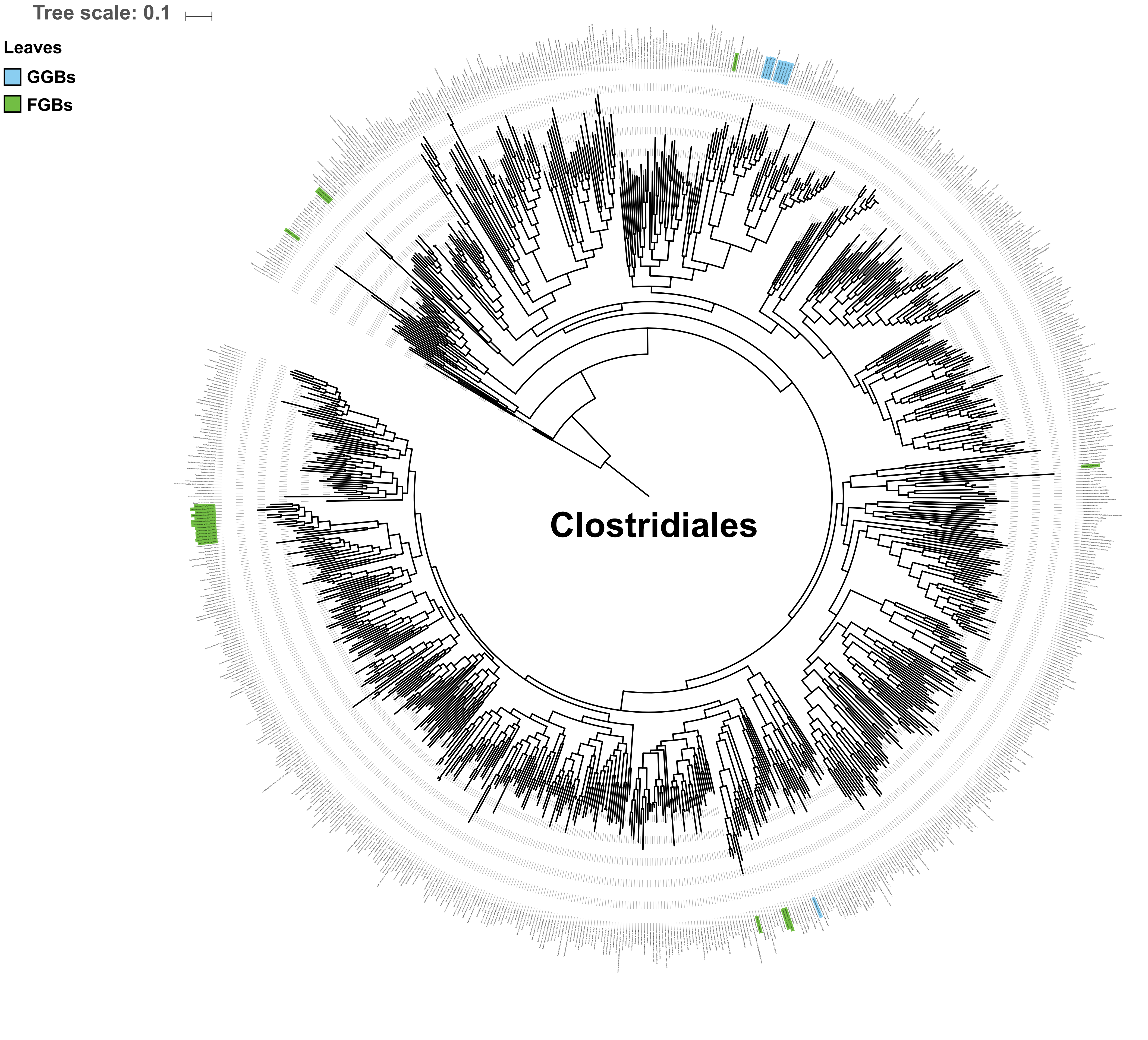

### Figure S8

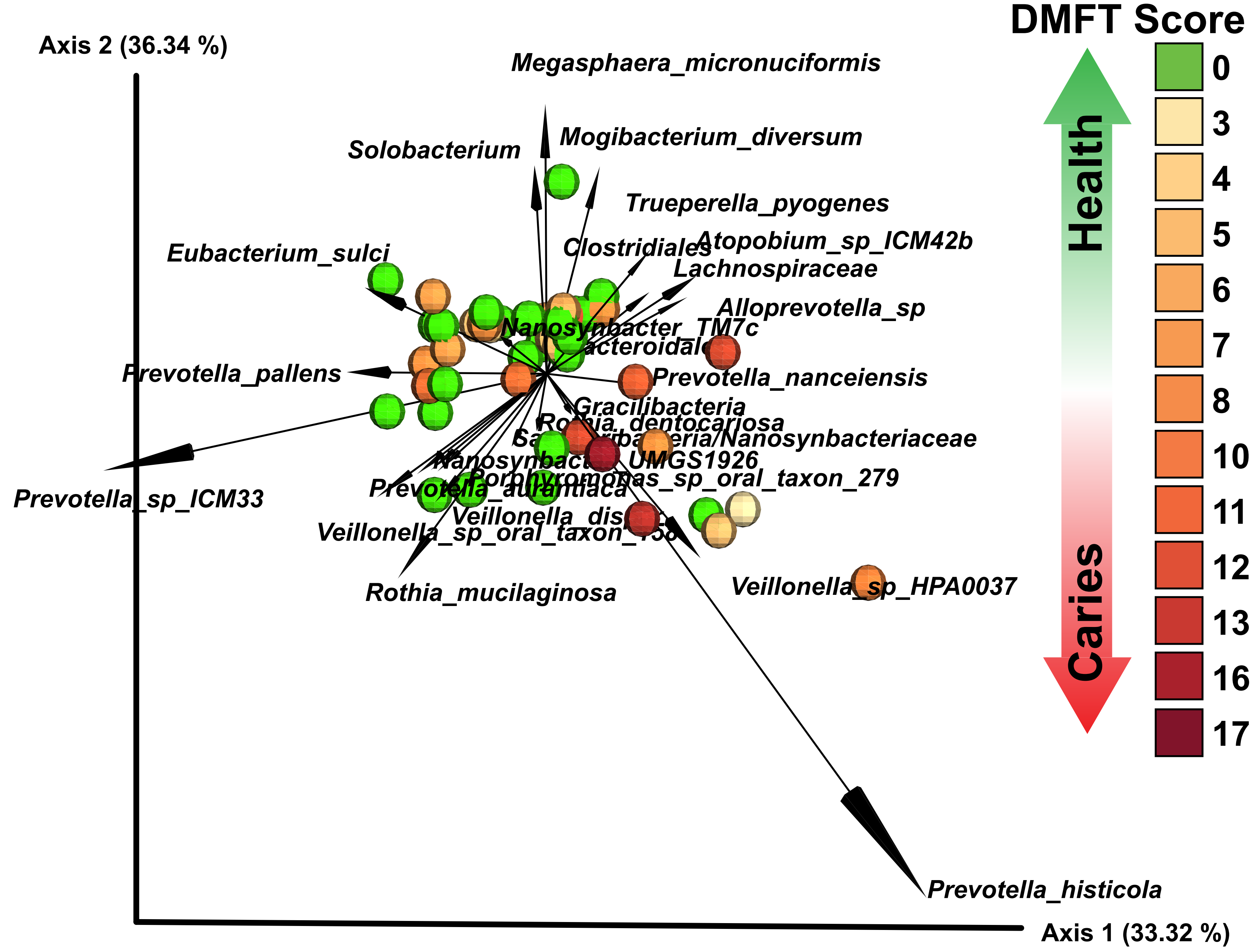

### Figure S9

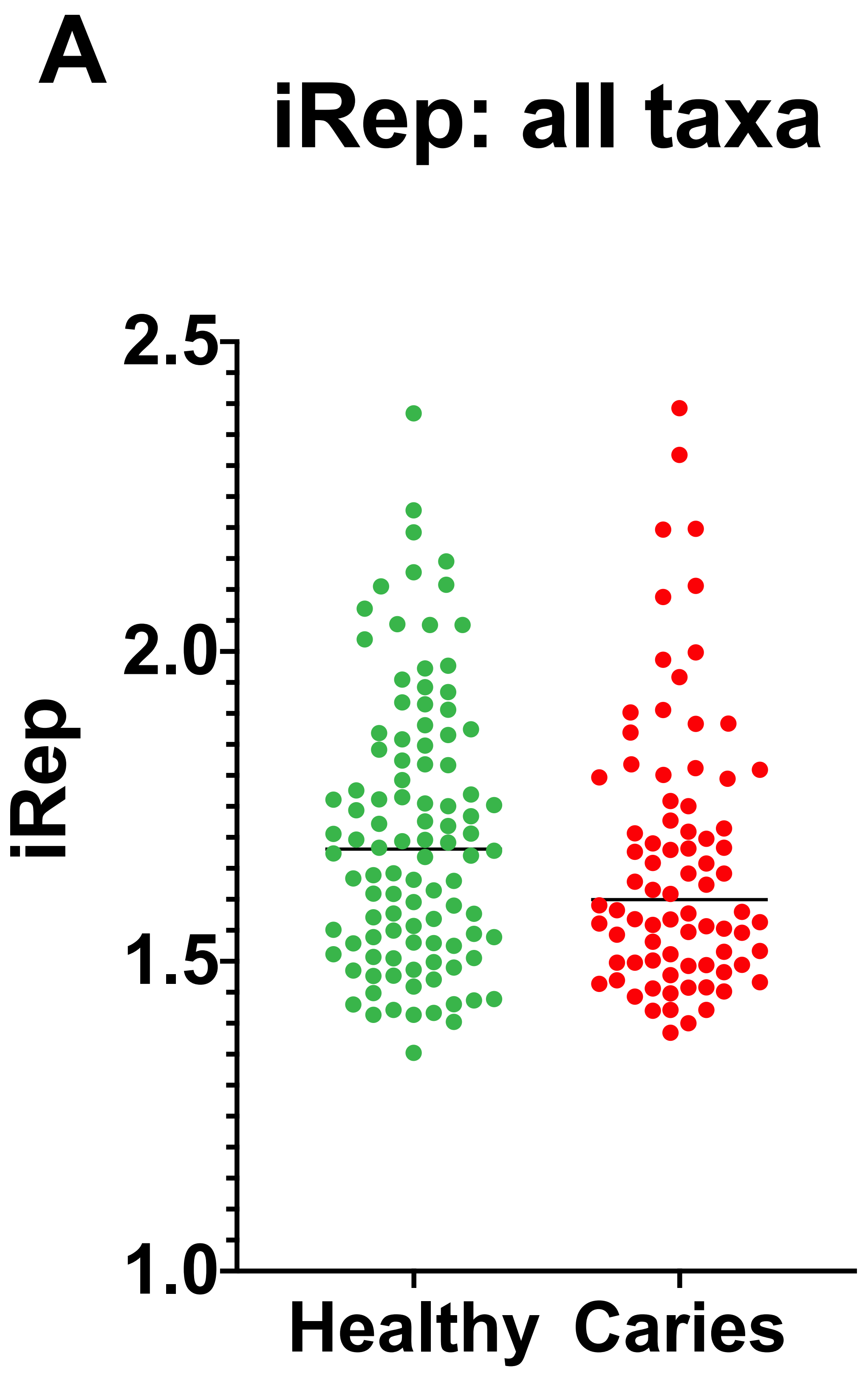

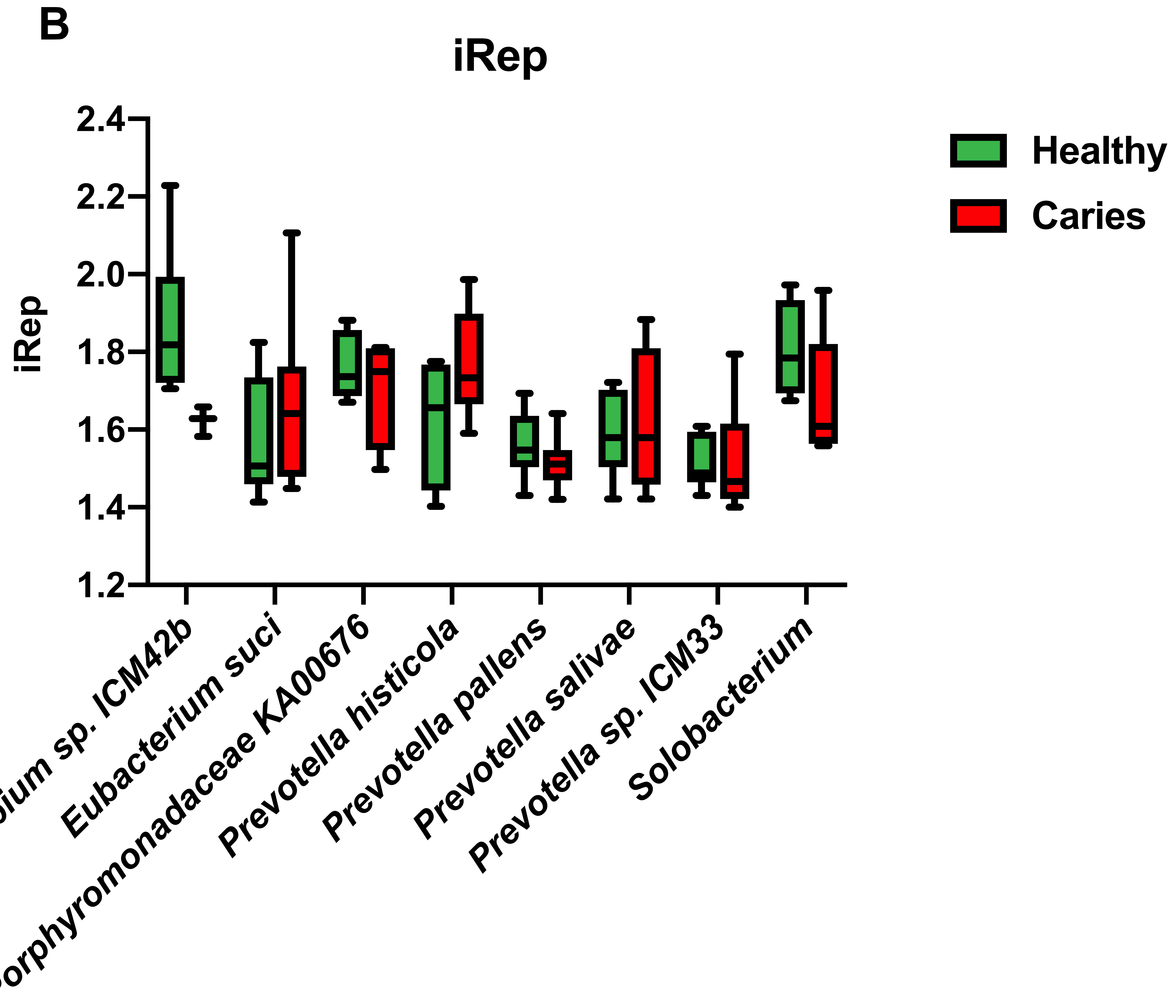
