## Supplemental Notes & Figure Legends for "Deep metagenomics examines the oral microbiome during dental caries, revealing novel taxa and co-occurrences with host molecules"

**Supplemental Appendix**

**Supplemental Notes**

**Supplemental Note 1. MAG assembly and binning yields 527 medium and high quality genomes, 20 representing novel species and 23 representing novel genera and species.** To uncover deeper details in the metagenomics dataset, and possibly identify novel taxa and genomic diversity, the quality-controlled Illumina sequencing reads were assembled using metaSPAdes (1). The resulting assemblies were binned and further improved using the MetaWRAP (2) pipeline. This pipeline has the advantage of utilizing an ensemble binning approach with three binning tools, Metabat2, Maxbin2, and Concoct. MetaWRAP then chooses the best bins from the three methods and reassembles these bins from the raw reads, providing significant improvements to the completeness, contamination, and N50 of the final bins (Dataset S2) (2). The completeness and contamination thresholds for all steps in MetaWRAP were >50% completeness and <10% contamination. This approach yielded 527 bins that were of at least Medium Quality according to the guidelines set forth by the Genomic Standards Consortium (GSC) regarding the Minimum Information about a Metagenome-Assembled Genome (MIMAG) (>50% completeness, <10% contamination)(Table S6) (3). Generally speaking, the samples with deeper sequencing provided more bins meeting this threshold, with sample SC33, with 249 million non-human reads yielding 69 bins, while sample SC26, with 1.4 million non-human reads yielding just 3 bins. A separate assembly was performed for each sample, as opposed to a co-assembly assembly of all samples, an alternative approach used by some studies. The pros and cons of a co-assembly versus individual assemblies have been discussed previously (4). As a result of the individual assemblies, many of the 527 MAGs were likely to represent redundant species across samples. To address this issue and dereplicate the MAGs, fastANI (5) was used to create an all vs. all % ANI distance matrix using a threshold of an average nucleotide identity (ANI) >95%, a threshold utilized by several recent landmark metagenomics studies to demark the ‘species’ unit (Figure 5A) (4-7). This process yielded 151 species-level genome bins (SGBs) (Figure 5A). To determine taxonomy, each of the 527 MAGs was compared to all genomes in the RefSeq database using Mash (8). All MAGs with a Mash distance of <5 (corresponding to a >95% ANI) to a RefSeq genome were annotated as the reference species with the shortest Mash distance. This process yielded 95 known SGBs (kSGBs), representing 376 MAGs and 56 unknown SGBs (uSGBs), representing 126 MAGs. In the vast majority of cases, all MAGs in each SGB had the same closest Mash hit, serving as a valuable sanity check of the ANI distance matrix. To further curate the uSGBs, each uSGB was examined for taxonomy using MetaWRAP (via taxator tk and Kraken) and blastx. Each uSGB was then compared against all genomes in the wider GenBank database under the predicted taxonomy using fastANI. Following this analysis, 15 uSGBs, representing 30 MAGs, were reassigned to kSGBs, as they had a >95% ANI match in GenBank (Table S7). There were 20 bins representing 50 MAGs that had a >85% and <95% ANI match to a GenBank genome. These were termed genus-level genome bins (GGBs), as the genus can be assigned with a fair amount of confidence, while the species appears to be not previously described. 23 bins, representing 48 MAGs had no match reference in GenBank with an ANI >85%. These were termed family-level genome bins (FGBs), as the family can be inferred, but the MAGs likely represent novel genera. Although the GGBs and FGBs on average had lower completion, higher contamination, a smaller N50, and more contigs/Mbp, the differences were not extensive, and the FGBs actually had a higher completeness and lower contigs/Mbp than GGBs (Figure 5B-E).

PhyloPhlAn2 (4) was used to phylogenetically place the uSGBs amongst reference strains at the order or class level. The PhyloPhlAn2 phylophlan_metagenomics.py script was also used on this set of MAGs and produced comparable results to the initial methods used here, and served as an independent validation of the approaches utilized to assemble and taxonomically classify the SGBs (Table S8). 25 of the MAGs, including 6 GGB and 11 FGB appeared to be Candidate Phyla Radiation (CPR) bacteria. This recently described supergroup is predicted to contain >35 phyla representing >15% of the diversity of all bacteria (9). CPR taxa have long been considered microbial “dark matter” and only one species has been cultivated thus far (10). CPR have reduced genomes and are thought to be obligate epibionts (11). In this MAGs dataset, 22 CPR MAGs were Saccharibacteria (formerly known as TM7), while 2 CPR MAGs were Gracilibacteria and one was an Absconditabacteria (formerly known as SR1). A comprehensive phylogeny of currently available Saccharibacteria was recently reported and novel named taxonomic hierarchies proposed (Jeff bioRXIV). In the present study, the Saccharibacteria MAGs represent Groups 1 (order Nanosynbacteriaceae), 3 (order Nanosynsoccalia), and 6 (order Nanoperiomorables). A table with information regarding the CPR MAGs reported here is provided in Table S9 and the phylogenetic trees are provided in Figure 4F (for Saccharibacteria) and Figure S5. 3 Saccharibacteria MAGs were the previously reported species TM7c (12). The Saccharibacteria MAG, 1_bin.14, had a 95% ANI to the previously reported species UMGS1926 (7), a Group 1 Saccharibacteria, 2 Saccharibacteria MAGs had a 95% ANI to a Saccharibacteria genome reported (13) and subsequently refined (14) previously, which appears to be in within the Saccharimonia class. The MAG 32_bin.49 was a species whose genome was also reported and refined in the same two studies (13, 14), which appears to be within the Nanogingivales order (TM7 G6). All other Saccharibacteria MAGs reported in this study appear to be novel genera and/or species. The Absconditabacteria MAG reported here has a 95% ANI to a previously isolated Absconditabacteria MAG (14). Of the two Gracilibacteria MAGs reported here, 1 had a 98% ANI to a previously reported genome (NCBI: NZ_CP017714.1), while the other appears to be a completely novel lineage.

Several of the uSGBs were in the class Bacteroidales. Among these, the a GGB containing 3 MAGs was found to be *Alloprevotella tannerae*. Two FGBs, representing 5 total MAGs, appear to be a more distant clade within the *Alloprevotella* branch. Another FGB, containing 5 MAGs, was phylogenetically placed within *Prevotella,* next to *Prevotella multisaccharivorax,* but is has only 74% ANI to the nearest reference taxa, *Prevotella oulorum.* The FGB/MAG 32_bin.69 was placed within the *Paludibacter* clade, while two FGBs (44_bin.40 and 44_bin.22) were placed within the *Porphyromonas* clade. The phylogenetic tree of the Bacteroidales uSGBs found in this study is provided in Figure S6.

A number of the uSGBs were within the highly polyphyletic order, Clostridiales. One FGB (47_bin.6 and 44_bin.29) was placed next to *Catonelli morbi,* which represented the highest %ANI to those MAGs, yet had only 77% ANI to that taxa. A notable FGB, containing 13 MAGs, had a highest %ANI (80%) with *Oribacterium,* yet was placed into the Clostridiales tree adjacent to *Sarcina* and *Butyrovibrio.* Clostridiales FGB 44_bin.37 was placed in the clade containing *Eubacterium brachy* and *Eubacterium saphenum.* The *Peptostreptococcus* GGB, containing 8 MAGs, was placed in a clade containing *Peptostreptococcus stomatis, Peptostreptococcus sp. D1,* and *Peptostreptococcus sp. MV1.* The FGB containing MAG 44_bin.10 was placed adjacent to the clade containing *Peptoanaerobacter stomatis.* Finally, 3 FGBs, containing 6 total MAGs, were placed in a distant area of the Clostridiales tree with few named neighbors. The phylogenetic tree of the Clostridiales uSGBs found in this study is provided in Figure S6.

Overall, the taxonomy of the assembled MAGs largely reflected the taxonomy of the communities predicted by MetaPhAn2. One notable exception was the lack of Streptococci among the assembled bins. Difficulty assembling quality Streptococcal genomes from metagenomics datasets has been noted previously (13), and is thought to occur because the high promiscuity of *Streptococcus* k-mers (due to high intra-genera diversity within Streptococcus, particularly from oral samples). Assembly biases such as this are examples that support use of unassembled reads for taxonomic abundance profiling and similar analyses. With this caveat in mind, the quantity of each assembled bin (MetaWRAP quant bins module) was used to create taxonomic abundance table, which was analyzed by DEICODE (Figure S8). Many of the overall trends were preserved compared to the DEICODE analysis of the unassembled reads, however the authors recommend utilizing unassembled reads for abundances purposes to avoid biases introduced by the assembly process.

**Supplemental Note 2. Estimation of the actively replicating taxa using iRep.** To estimate which taxa identified in the metagenomics analysis were alive and metabolically active, the bioinformatics tool iRep was utilized (15). iRep infers replication rates based upon differential sequencing coverage of genomic regions with respect to the origin of replication (15). iRep requires 75% completeness, <175 contigs per 1mb, and looks at contigs > 5000bp with a > 5 average coverage. iRep was able to compute replication rates for 183 of 527 MAGs. The replication rate of all bacteria was not significantly different among the taxa derived from the saliva of healthy children compared to the saliva from the children with caries (Figure S9A). At the individual SGB level, there were only 8 SGBs with sufficient (at least 3) MAGs, which passed the requirements for iRep, from both the healthy and the caries groups. None of these 8 SGBs had statistically different rates of replication between the healthy and caries groups (Figure S9B).

The iRep tool provides a unique and highly useful way to examine the replication rate (and therefore “activity” or “viability”) of bacteria given only genomic DNA or MAGs (15). The caveat is the requirement of genomes or MAGs to meet several criteria of completeness and fragmentation. These constrains limited the number of genomes this study was able to analyze using iRep, particularly when sufficient MAGs from both caries and healthy study groups were needed for comparison. Although there were no statistically different replication rates detected between the caries and healthy groups in this study, more extensive application of this tool would be very likely to provide interesting information regarding the oral microbiome.

**Supplemental Figure Legends**

**Fig S1. Heatmap illustrating species-level taxonomic abundances in the caries versus healthy groups.** Plot was generated using the ComplexHeatmaps R package (16). Abundances were determined using MetaPhAn2 (17). Rows and columns were clustered using a Bray-Curtis distance matrix and the “complete” clustering method. Only taxa appearing in at least 10 samples are included in this figure.

**Fig S2. Heatmap illustrating species-level functional pathway abundances in the caries versus healthy groups.** Plot was generated using the ComplexHeatmaps R package (16). Abundances were determined using HumanN2 (18). Rows and columns were clustered using a Bray-Curtis distance matrix and the “complete” clustering method. Only pathways included in at least 10 samples are included in this figure.

**Fig S3. Plot illustrating inverse Pearson correlation between number of caries lesions and alpha diversity (Shannon Index) of functional pathways.** Plot generated using the QIIME 2 (19) diversity plugin.

**Fig S4. Low dimensionality of functional pathways makes pathway-host marker co-occurrences difficult to identify and interpret.** Biplot generated using QIIME 2 (19, 20), co-occurrence vectors were calculated using MMVEC. Spheres indicate host markers, which are colored according to whether each was significantly increased in the caries group (red) or not (blue). Functional pathways are represented by the vectors (arrows), which are colored according to the differential rank of each pathway with respect to disease status as determined by Songbird (21).

**Fig S5. Phylogenetic placement of unknown species-level genome bins (uSGBs) within Candidate Phyla Radiation (CPR).** Trees were generated using PhyloPhlAn2 (4) and visualized using iToL (22). **(A) Saccharibacteria (formerly TM7)**. Tree leaves are highlighted to illustrate MAGs assembled in this study and highlight is colored by bin type (SGBs = orange; GGBs = blue; FGBs = green;*Nanosynbacter lyticus TM7x* = yellow). Branches are colored by TM7 group (as proposed by McLean et al.). **(B) Gracilibacteria** and **(C) Absconditabacteria.**

**Fig S6. Phylogenetic placement of uSGBs within Bacteroidales order.** Tree was generated using PhyloPhlAn2 and visualized using iToL. Tree leaves are highlighted to illustrate MAGs assembled in this study and highlight is colored by bin type (SGBs [reassigned using GenBank] = orange; GGBs = blue; FGBs = green).

**Fig S7. Phylogenetic placement of uSGBs within Clostridiales class.** Tree was generated using PhyloPhlAn2 and visualized using iToL. Tree leaves are highlighted to illustrate MAGs assembled in this study and highlight is colored by bin type (= orange; GGBs = blue; FGBs = green).

**Fig S8. Beta diversity of taxa using quantification of assembled genome bins.** 3D PCA plot generated using DEICODE (robust Aitchison PCA) (23). Data points represent individual subjects and are colored with a gradient to visualize DMFT score, indicating severity of dental caries. Feature loadings (i.e. taxa driving differences in ordination space) are illustrated by the vectors, which are labeled with the cognate species name. Taxonomic abundances were determined using the Quant Bins module of MetaWRAP (2).

**Fig S9. Replication rates of all taxa (A) and most abundant taxonomic groups (B).** Replication rates were determined using iRep (15). Replication rates were not significantly different between healthy and caries groups for any taxonomic groups shown here for overall (A).

**Supplemental References**

1.	Nurk S, Meleshko D, Korobeynikov A, Pevzner PA. 2017. metaSPAdes: a new versatile metagenomic assembler. Genome Res 27:824-834.

2.	Uritskiy GV, DiRuggiero J, Taylor J. 2018. MetaWRAP-a flexible pipeline for genome-resolved metagenomic data analysis. Microbiome 6:158.

11.	Baker JL, Bor B, Agnello M, Shi W, He X. 2017. Ecology of the Oral Microbiome: Beyond Bacteria. Trends Microbiol 25:362-374.

12.	Marcy Y, Ouverney C, Bik EM, Losekann T, Ivanova N, Martin HG, Szeto E, Platt D, Hugenholtz P, Relman DA, Quake SR. 2007. Dissecting biological "dark matter" with single-cell genetic analysis of rare and uncultivated TM7 microbes from the human mouth. Proc Natl Acad Sci U S A 104:11889-94.

13.	Espinoza JL, Harkins DM, Torralba M, Gomez A, Highlander SK, Jones MB, Leong P, Saffery R, Bockmann M, Kuelbs C, Inman JM, Hughes T, Craig JM, Nelson KE, Dupont CL. 2018. Supragingival Plaque Microbiome Ecology and Functional Potential in the Context of Health and Disease. MBio 9.

14.	Shaiber A, Eren AM. 2019. Composite Metagenome-Assembled Genomes Reduce the Quality of Public Genome Repositories. MBio 10.

15.	Brown CT, Olm MR, Thomas BC, Banfield JF. 2016. Measurement of bacterial replication rates in microbial communities. Nat Biotechnol 34:1256-1263.

16.	Gu Z, Eils R, Schlesner M. 2016. Complex heatmaps reveal patterns and correlations in multidimensional genomic data. Bioinformatics 32:2847-9.

17.	Truong DT, Franzosa EA, Tickle TL, Scholz M, Weingart G, Pasolli E, Tett A, Huttenhower C, Segata N. 2015. MetaPhlAn2 for enhanced metagenomic taxonomic profiling. Nat Methods 12:902-3.

18.	Franzosa EA, McIver LJ, Rahnavard G, Thompson LR, Schirmer M, Weingart G, Lipson KS, Knight R, Caporaso JG, Segata N, Huttenhower C. 2018. Species-level functional profiling of metagenomes and metatranscriptomes. Nat Methods 15:962-968.

19.	Bolyen E, Rideout JR, Dillon MR, Bokulich NA, Abnet CC, Al-Ghalith GA, Alexander H, Alm EJ, Arumugam M, Asnicar F, Bai Y, Bisanz JE, Bittinger K, Brejnrod A, Brislawn CJ, Brown CT, Callahan BJ, Caraballo-Rodriguez AM, Chase J, Cope EK, Da Silva R, Diener C, Dorrestein PC, Douglas GM, Durall DM, Duvallet C, Edwardson CF, Ernst M, Estaki M, Fouquier J, Gauglitz JM, Gibbons SM, Gibson DL, Gonzalez A, Gorlick K, Guo J, Hillmann B, Holmes S, Holste H, Huttenhower C, Huttley GA, Janssen S, Jarmusch AK, Jiang L, Kaehler BD, Kang KB, Keefe CR, Keim P, Kelley ST, Knights D, et al. 2019. Reproducible, interactive, scalable and extensible microbiome data science using QIIME 2. Nat Biotechnol 37:852-857.

20.	Vazquez-Baeza Y, Pirrung M, Gonzalez A, Knight R. 2013. EMPeror: a tool for visualizing high-throughput microbial community data. Gigascience 2:16.

21.	Morton JT, Marotz C, Washburne A, Silverman J, Zaramela LS, Edlund A, Zengler K, Knight R. 2019. Establishing microbial composition measurement standards with reference frames. Nat Commun 10:2719.

22.	Letunic I, Bork P. 2019. Interactive Tree Of Life (iTOL) v4: recent updates and new developments. Nucleic Acids Res 47:W256-W259.

23.	Martino C, Morton JT, Marotz CA, Thompson LR, Tripathi A, Knight R, Zengler K. 2019. A Novel Sparse Compositional Technique Reveals Microbial Perturbations. mSystems 4.
